# The Microbial Metabolite 4-Cresol Improves Glucose Homeostasis and Enhances β-Cell Function

**DOI:** 10.1101/444893

**Authors:** Francois Brial, Fawaz Alzaid, Kazuhiro Sonomura, Kelly Meneyrol, Aurélie Le Lay, Noémie Péan, Ana Neves, Julien Chilloux, Lyamine Hedjazi, Taka-Aki Sato, Andrea Rodriguez Martinez, Nicolas Venteclef, Jeremy K Nicholson, Christophe Magnan, Mark Lathrop, Marc-Emmanuel Dumas, Fumihiko Matsuda, Pierre Zalloua, Dominique Gauguier

**Affiliations:** Sorbonne University, University Paris Descartes, INSERM UMR_S 1138, Cordeliers Research Centre, 75006 Paris, France; Center for Genomic Medicine, Kyoto University Graduate School of Medicine, Kyoto 606-8501, Japan; Life Science Research Center, Technology Research Laboratory, Shimadzu Corporation, Kyoto 604-8511, Japan; Sorbonne Paris Cité, University Denis Diderot, CNRS UMR 8251, 75205 Paris, France; Imperial College London, Section of Biomolecular Medicine, Division of Computational and Systems Medicine, Department of Surgery and Cancer, Faculty of Medicine, Sir Alexander Fleming building, London SW7 2AZ, United Kingdom; McGill University and Genome Quebec Innovation Centre, 740 Doctor Penfield Avenue, Montreal, QC, H3A 0G1, Canada; Lebanese American University, School of Medicine, Beirut, 1102 2801 Lebanon.

**Keywords:** Metabolome, Gut Microbiome, Type 2 Diabetes, Obesity, Insulin Secretion, Pancreatic Islets, Beta-cells, Metabotype, Mouse, Rat

## Abstract

Gut microbiota changes are associated with increased risk of Type 2 diabetes (T2D) and obesity. Through serum metabolome profiling in patients with cardiometabolic disease (CMD) we identified significant inverse correlation between the microbial metabolite 4-cresol and T2D. Chronic administration of non toxic dose of 4-cresol in two complementary preclinical models of CMD reduced adiposity, glucose intolerance and liver triglycerides, and enhanced insulin secretion *in vivo*, which may be explained by markedly increased pancreas weight, augmented islet density and size, and enhanced vascularisation suggesting activated islet neogenesis. Incubation of isolated islets with 4-cresol enhanced insulin secretion, insulin content and cell proliferation. In both CMD models 4-cresol treatment *in vivo* was associated with altered expression of SIRT1 and the kinase DYRK1A, which may contribute to mediate its biological effects. Our findings identify 4-cresol as an effective regulator of β-cell function and T2D endophenotypes, which opens therapeutic perspectives in syndromes of insulin deficiency.

---

Type 2 diabetes (T2D) and associated elements of cardiometabolic diseases (CMD) are multifactorial disorders resulting from the effects of many genes interacting with environmental factors. The gut microbiota sits at the interface between the environment and the host organism and contributes to the metabolism of nutrients into digestible molecules and the maturation of immune processes. The involvement of the gut microbiota in T2D and obesity has been mainly addressed through metagenomic sequencing, which identified changes in microbiome architecture ^1-3^ and gene richness ^4^. This concept of microbiome-host crosstalk is also supported by instances of associations between CMD and microbial-mammalian co-metabolites (eg. hippurate, methylamines, short chain fatty acids) ^5^ that metabolomic technologies can detect and quantify in biofluids. Metabolomics is able to quantitatively and qualitatively analyze a broad range of small molecules, which are molecular endpoints of the combined expression of the host genome and the microbiome ^6^. Metabolic endophenotypes generated by metabolomic-based phenotyping and metabotyping ^7^ can be used as biomarkers underlying disease etiology ^8,9^ and help understand disease pathogenesis and develop relevant therapeutic solutions.

Considering the importance of microbiome-host metabolic axis in chronic diseases ^10,11^, we set out to investigate interactions of risk factors for T2D and obesity and gut microbial metabolites. We used a semi-targeted gas chromatography coupled to mass spectrometry (GC-MS) approach to profile plasma metabolites and test their association with T2D and obesity. Simultaneous analysis of 101 metabolites identified a metabotype of 18 metabolites associated with the diseases, including products of gut microbial metabolism (eg. 4-cresol). Chronic administration of 4-cresol in preclinical models of spontaneous or experimentally-induced insulin resistance resulted in significantly improved glucose homeostasis, enhanced glucose-stimulated insulin secretion *in vivo*, reduced adiposity and increased pancreas weight. Further physiological, histological and molecular analyses in 4-cresol treated animals indicated reduction in inflammation and enhancement of both cell proliferation and vascularisation in the pancreas, which may be mediated by downregulated expression of the kinase DYRK1A by 4-cresol ^12^ and upregulated expression of SIRT1. Our findings illustrate the contribution of gut microbial metabolites in chronic diseases through the identification of 4-cresol as a regulator of T2D endophenotypes with potential therapeutic applications.

## RESULTS

### Clinical and biochemical data analysis

The study population consisted of 138 phenotypically well characterized patients with a mean age of 54.96 (± 1.01) years, a mean body weight of 77.84 kg (± 1.23) and a mean BMI of 28.04 kg/m^2^ (± 0.42). T2D identification was based on clinical documentation in the patients’ medical charts. Among these, 77 were classified as high CMD risk patients with elevated BMI (>30kg/m^2^) (24%), low plasma HDL cholesterol (<40mg/dL) (50%) and fasting hyperglycemia (>125mg/dL) (10%). These were classified as cases (77), whilst the remaining 61 subjects were classified as control. There were no significant gender differences for any of these phenotypes.

### Metabolic profiling identifies a metabotype associated with T2D and obesity

To identify quantitative variations in circulating metabolites that can account for clinical features and biochemical measures in the population, serum samples from cases and controls were processed by GC-MS for metabolic profiling. A total of 101 metabolites were targeted in the GC-MS spectra and used for association analysis with clinical phenotypes in the study population (**Supplementary Table 1**). The targeted metabolites included 36 amino acids, 32 organic acids, 20 sugars, and other molecular compounds (eg. nucleobases).

We carried out focused analyses of association between metabolites and T2D and obesity. We first built an orthogonal partial least squares discriminant analysis (O-PLS-DA) using GC-MS data to stratify the population based on T2D and obesity (**Figure 1A**). The goodness-of-prediction (Q^2^_Yhat_ statistic = 0.36) of the model assessed by 7-fold cross-validation was deemed significant by permutation testing (p = 10^-5^ based on 100,000 iterations) (**Figure 1B**). Model coefficients supporting this discrimination were validated against logistic regressions adjusted for age and gender and highlighted a signature made of 14 metabolites (2-aminoadipic acid, arabinose, 2-deoxytetronic acid, gluconic acid, glucose, 2-hydroxybutyric acid, 3-hydroxyisobutyric acid, 2-and 3-hydroxyisovaleric acid, indolelactic acid, isocitric acid, mannose, ribulose, valine) significantly positively correlated with T2D and obesity and 4 metabolites (1,5-anhydro-D-sorbitol, 4-cresol, glyceric acid and 5-oxoproline) negatively correlated with the diseases (**Figure 1C**). 4-cresol was the microbial metabolite the most significantly associated with reduced disease risk.

**Figure 1.**
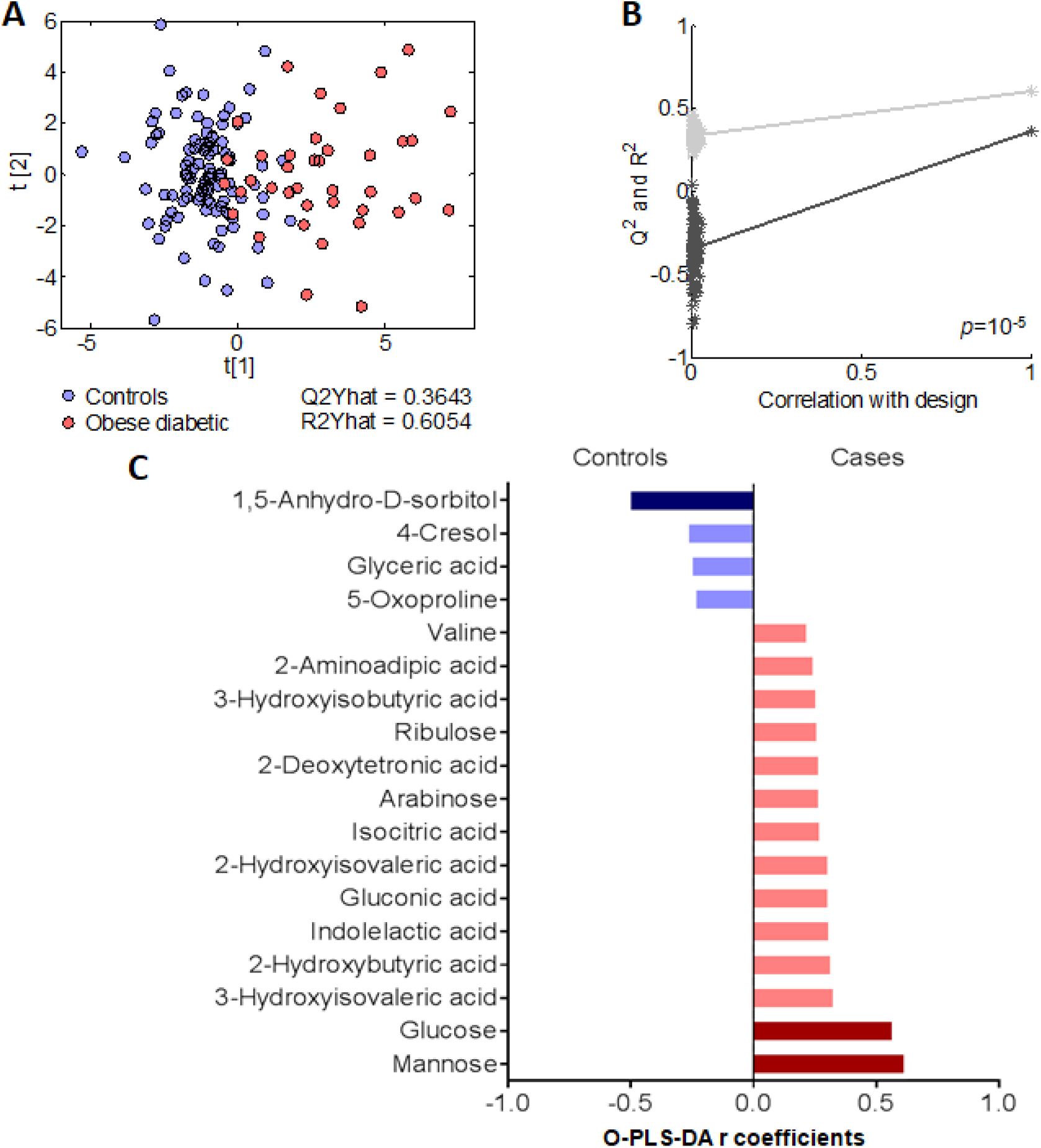
Metabolic profiling identifies microbial metabolites associated with type 2 diabetes. Data from serum metabolites acquired by gas chromatography mass spectrometry (GC-MS) and clinical variables in the cohort were used to derive OPLS-DA scores plot (A). Validation of goodness-of-fit (R^2^) and goodness-of-prediction (Q^2^) parameters were obtained by permutation testing (n=100 random permutations) (B). Variable contributions (r) to the O-PLS-DA model, confirmed by logistic regression are shown in green and red for negative and positive correlations with the disease, respectively. Statistics were corrected for multiple comparisons and adjusted for age and gender effects.

These results demonstrate the power of systematic and simultaneous metabolomic-based profiling of a large series of metabolites to identify metabotypic endophenotypes associated with diseases, and point to the microbial metabolite 4-cresol as a novel biomarker of reduced risk of T2D and obesity.

### 4-cresol treatment improves glucose homeostasis and reduces body weight gain in mice

To test the biological role of microbial metabolites *in vivo* we prioritized experimental validation of the clinical association on 4-cresol, which suggested potential benefits for T2D and obesity. We chronically treated mice fed control diet or high fat diet (HFD) with subcutaneous infusion of a solution of 4-cresol 0.04M, corresponding to approximately 0.5mg/kg/day, for 42 days. Subcutaneous treatment was preferred over dietary supplementation or oral gavage in order to control permanent delivery of 4-cresol over a long period of time and reduce the stress induced by animal handling. This delivery method also avoided possible toxic effects of this volatile compound that are observed at much higher doses (240-2000mg/kg/day) on neurological function, liver function and respiratory epithelium ^13^. As expected, mice fed HFD rapidly gained more weight than mice fed control diet and developed fasting hyperglycemia and marked glucose intolerance (**Figure 2A-F**). Glycemia after the glucose challenge, cumulative glycemia during the test and the ΔG parameter were significantly more elevated in mice fed HFD than in controls (**Figure 2C-F**). Fasting insulin and glucose-induced insulin secretion were not significantly affected by HFD (**Figure 2G**).

**Figure 2.**
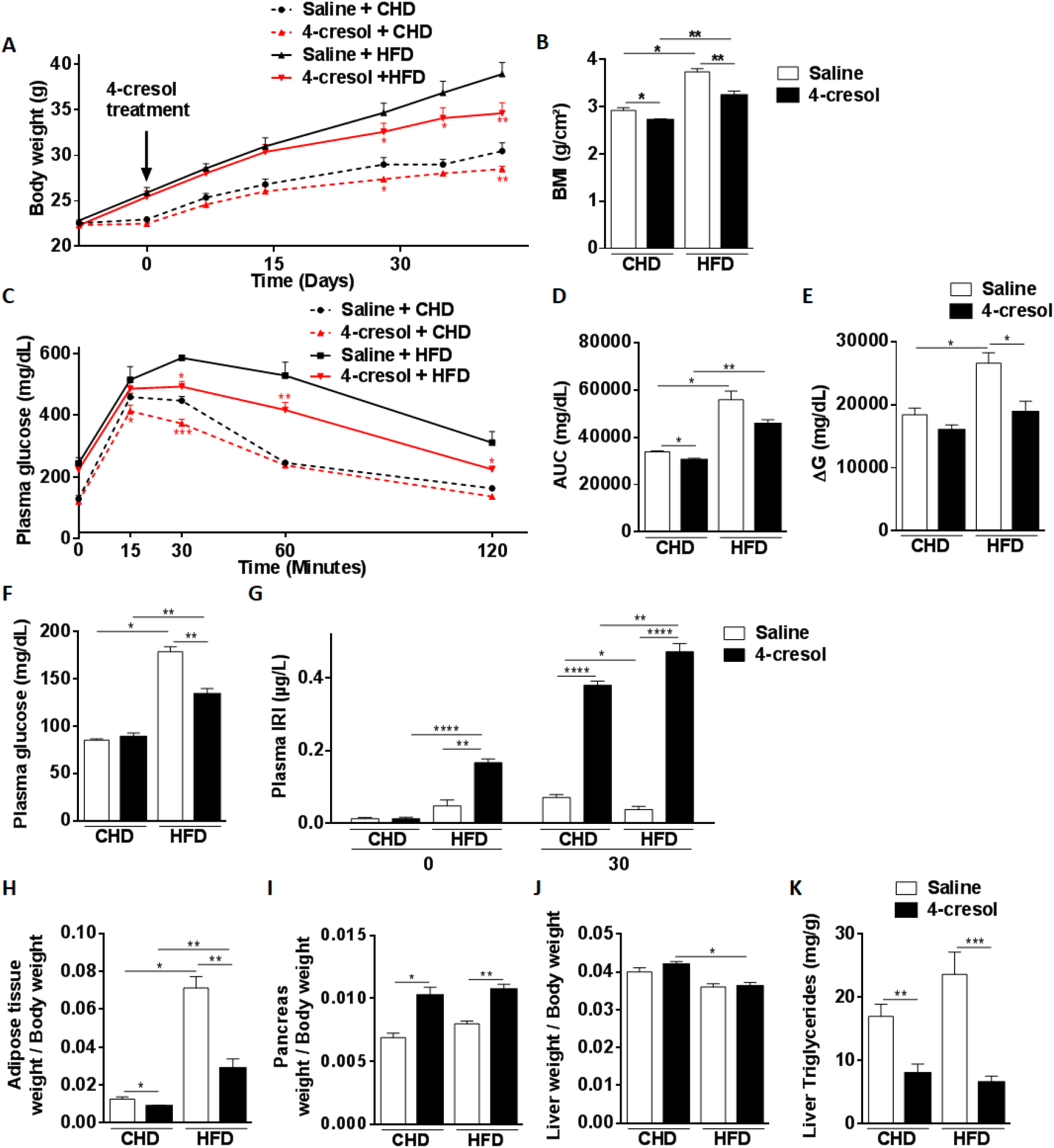
Chronic administration of 4-cresol improves glucose homeostasis, stimulates insulin secretion and reduces obesity in C57BL6/J mice. The effects of 6-week long administration of 4-cresol *in vivo* in mice fed control chow or high fat diet (HFD) were tested on body weight (A), body mass index (BMI) (B), glucose homeostasis (C-F), glucose-stimulated insulin secretion (G), organ weight (H-J) and liver triglycerides (K). BMI was calculated as body weight divided by the squared of anal-nasal length. AUC was calculated as the sum of plasma glucose values during the IPGTT. ΔG is the AUC over the baseline value integrated over the 120 minutes of the test. All measures are from 6 mice per group. Data were analyzed using the unpaired Mann-Whitney test. Results are means ± SEM. *P < 0.05; **P < 0.01; ***P < 0.001; ****P < 0.0001, significantly different to relevant controls. IRI, immunoreactive insulin

Chronic infusion of mice fed control diet with 4-cresol resulted in progressive reduction in body weight when compared to mice treated with saline (**Figure 2A**). This effect became significant after 4 weeks of 4-cresol treatment and remained significant until the end of the experiment. BMI was also significantly decreased after six week of 4-cresol treatment (**Figure 2B**). Glucose tolerance was improved by 4-cresol, as indicated by the significant reduction in both acute glycemic response to the glucose challenge and cumulative glycemia during the IPGTT when compared to saline-treated mice (**Figure 2C,D**). 4-cresol infusion resulted in a significant increase in glucose-stimulated insulin secretion when compared to controls (**Figure 2G**).

The effect of 4-cresol on body weight, glycemic control and insulin secretion was also strongly significant in HFD-fed mice (**Figure 2A-G**). In addition, 4-cresol treatment in fat fed mice induced strong reduction in fasting glycemia (**Figure 2F**) and in the ΔG parameter, which was normalized to the level of mice fed chow diet and treated with saline (**Figure 2E**), and fasting hyperinsulinemia (**Figure 2G**) when compared to HFD-fed mice infused with saline.

### *In vivo* 4-cresol treatment alters organ weight and reduces liver triglycerides

To further characterize the effects of 4-cresol on phenotypes relevant to T2D and obesity, we measured organ and tissue weights, and analysed phenotypes frequently associated with obesity (eg. fatty liver disease) in the mouse groups. After seven weeks of HFD, mice exhibited significantly elevated adiposity, which was calculated as the ratio of the retroperitoneal fat pad weight to body weight (**Figure 2H**) and reduced heart weight (**Figure S1**) when compared to mice fed chow diet. Liver triglycerides levels were markedly increased by HFD but this effect was not statistically significant (**Figure 2K**).

In both diet groups, chronic administration of 4-cresol over 6 weeks resulted in significant reduction in adiposity index (**Figure 2H**) and increased pancreas weight by 48.9% (**Figure 2I**) when compared to mice treated with saline. Heart and kidney weight were not affected by 4-cresol (**Figure S1**). Liver weight was reduced in HFD-fed mice treated with 4-cresol (**Figure 2J**). 4-cresol treatment resulted in a systematic and dramatic reduction in liver triglycerides content in both diet groups (**Figure 2K**).

Collectively, these data demonstrate the beneficial effects of chronic administration of 4-cresol *in vivo* in mice on obesity, thus validating our results from metabolomic profiling in patients, and extend the characterization of its beneficial effects on improved glucose tolerance and reduced triglycerides in the liver.

### 4-cresol derivative 4-methylcatechol mimics the biological effects of 4-cresol

We then investigated whether these physiological effects are specific to 4-cresol or can be replicated with other products of microbial metabolism structurally related to 4-cresol. We repeated the *in vivo* phenotypic screening in control and HFD-fed mice treated with 4-methylcatechol (4-MC), a bacterial product of 4-cresol through enzymatic reactions potentially involving mono-oxygenase, dioxygenase and cycloisomerase ^14^. Chronic treatment with 4-MC in mice fed chow diet or HFD induced a strong reduction in body growth and BMI (**Figure S2A,B**), significant improvement in glycemic control on both diets, as demonstrated by improved glucose tolerance (**Figure S2C**), reduced cumulative glycemia and ΔG during the IPGTT (**Figure S2D,E**) and reduced fasting glycemia (**Figure S2F**), and increased insulin secretion (**Figure S2G**) when compared to mice treated with saline. Treatment of mice fed control diet or HFD with 4-MC induced significant reduction in adiposity, liver weight and liver triglycerides and increased pancreas weight (**Figure S2H-K**). These results indicate that 4-cresol and 4-MC have very similar effects in several tissues, and might regulate the same biological mechanisms in these tissues.

### Chronic 4-cresol treatment ameliorates histological features in adipose tissue and liver

To characterize the effects of 4-cresol on adiposity and lipid metabolism at the organ level, we carried out histological analyses of adipose tissue and liver in the four mouse groups (**Figure 3A-D**). HFD feeding induced a strongly significant increase (84.8%) in adipocyte size (56.27 ± 0.47 in CHD fed mice and 104.00 ± 1,41 in HFD fed mice) (**Figure 3A,B**), which is consistent with increased adiposity in response to fat feeding. In CHD fed mice 4-cresol induced a slight reduction in adipocyte diameter when compared to saline-treated mice (53.97 ± 0.39, P=0.006). Remarkably, 4-cresol administration in HFD fed mice led to a strongly significant reduction of adipocyte size (66.22 ± 0.70, P<0.0001) to a level close to that of saline treated CHD-fed controls (**Figure 3A,B**).

**Figure 3.**
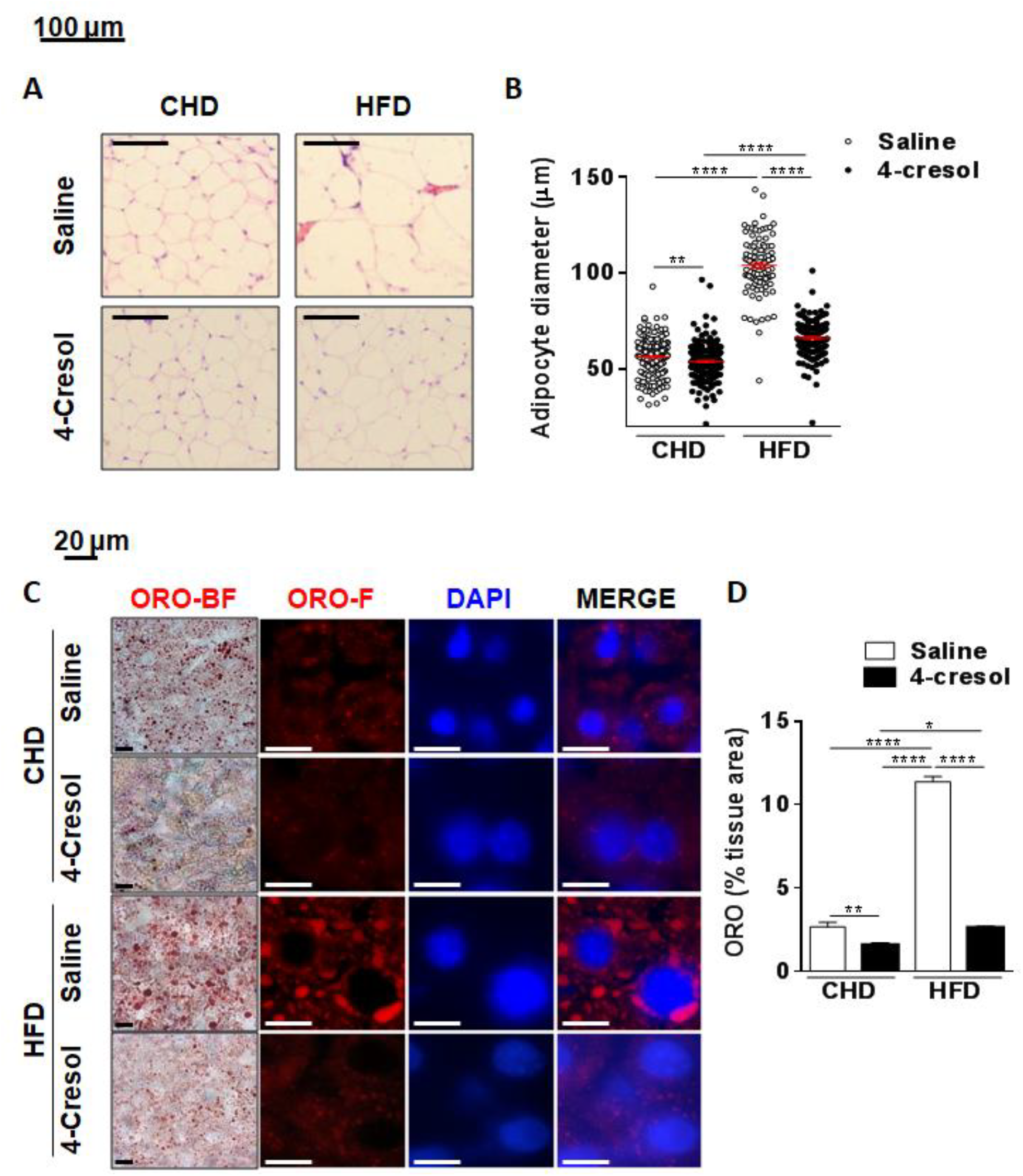
Chronic 4-cresol treatment ameliorates histological features in adipose tissue (A,B) and liver (C,D) of C57BL6/J mice. Mice were fed normal chow diet (CHD) or high fat diet (HFD) and treated with chronic infusion of either 4-cresol or saline for 6-weeks. Adipose tissue sections were labeled either with Hematoxylin-Eosin (HE) (A) to determine adipocyte size (B). Liver sections were labeled with Oil Red O (ORO) (C) to determine neutral fat content (D). All measures are from 6 mice per group. A total of 2000 cells (8000 cells per group) were analysed to determine adipocyte diameter. Data were analyzed using the unpaired Mann-Whitney test. Results are means ± SEM. *P<0.05; **P<0.01; ***P<0.001, significantly different to relevant controls. ORO-BF, ORO bright field; ORO-F, ORO fluorescent.

Occurrence of liver structural lesions resembling non alcoholic fatty liver disease (NAFLD) is a phenotypic hallmark consistently observed in obese mice fed HFD in our experimental conditions ^15^. To investigate the effect of 4-cresol on these defects and test the histological relevance of reduced liver triglycerides observed in mice treated by 4-cresol (**Figure 2K**), we carried out liver histology in the four mouse groups. HFD feeding induced a strongly significant 4.27-fold increase in liver fat content determined by Oil-Red-O staining of liver sections (**Figure 3C,D**). In response to 4-cresol infusion, liver fat content was reduced in mice fed CHD. It was also strongly reduced by 40.7% in HFD fed mice (P=0.009) to levels similar to saline-treated CHD-fed controls (**Figure 3D**).

### Gene expression changes induced by chronic 4-cresol treatment in adipose tissue

To investigate molecular changes potentially underlying morphological changes in adipose tissue caused by 4-cresol treatment, we analysed the expression of a selection of genes known to regulate adipocyte function in obesity (**Figure S3**). Expressions of sirtuin 1 (Sirt1), caveolin 2 (Cav2), hormone-sensitive lipase (Hsl) and patatin-like phospholipase domain containing 2 (Pnpla2, Atgl) were significantly stimulated by 4-cresol in CHD fed mice, whereas uncoupling protein 1 (Ucp1) expression was markedly downregulated. Expression of Hsl and Pnpla2 remained significantly upregulated by 4-cresol in HFD-fed mice when compared to saline treated fat fed mice, and Ucp1 was significantly downregulated in these mice when compared to both HFD-fed mice treated with saline and CHD-fed mice treated with 4-cresol.

### Chronic 4-cresol administration increases pancreatic vascularisation and islet density and promotes cell proliferation

One of our most striking observations was the massive effect of 4-cresol on increased pancreas weight in both diet groups (**Figure 2I**), which we analysed further through histology of pancreas sections. To test whether islet structure was affected by 4-cresol, we focused histopathology analyses on islets in sections stained by HE. Fat feeding did not affect overall insulin positive area (58.95 ± 11.75 in CHD fed mice; 109.03 ± 24.15 in HFD fed mice, P=0.071) (**Figure 4A,B**), but significantly increased islet density (**Figure 4C,D**). 4-cresol administration was associated with a strong increase in both islet density and insulin positive area in mice fed CHD (185.40 ± 22.95) or HFD (171.90 ± 21.62), but the effect was statistically significant only in CHD-fed mice (**Figure 4A-D**). We noted that islets in pancreas sections of cresol treated mice were predominantly located in the close vicinity of the vasculature (**Figure 4G**), suggesting an effect of 4-cresol on enhanced islet neogenesis. To investigate the possible cause of increased β-cell area and islet density induced by 4-cresol, we next used Ki67 to determine pancreatic cell proliferation in the four mouse groups (**Figure 5A-C**). Immunohistochemistry confirmed elevated islet size 4-cresol treated mice (153.60 ± 14.18 in CHD fed mice; 170.00 ± 12.56 in HFD fed mice) when compared to CHD-fed mice treated with saline (50.81 ± 2.90) (**Figure 5A,B**). The number of proliferative nuclei was increased in HFD-fed mice when compared to CHD-fed mice (**Figure 5A-C**). It was also significantly increased in response to 4-cresol treatment in CHD-fed mice and remained elevated in HFD-fed mice. Increased vascularisation from endothelial cells contributes to pancreatic cell proliferation and islet neogenesis, and may explain the effect of 4-cresol on cell proliferation. To test this hypothesis, we then stained pancreas sections of the four mouse groups with CD31, a marker of vascularisation. The number of CD31 positive cells was significantly increased in response to HFD (**Figure 5D,E**). In both groups fed CHD or HFD, 4-cresol induced a further significant increase in CD31 positive cells, thus demonstrating the role of this metabolite on the stimulation of pancreas vascularisation.

**Figure 4.**
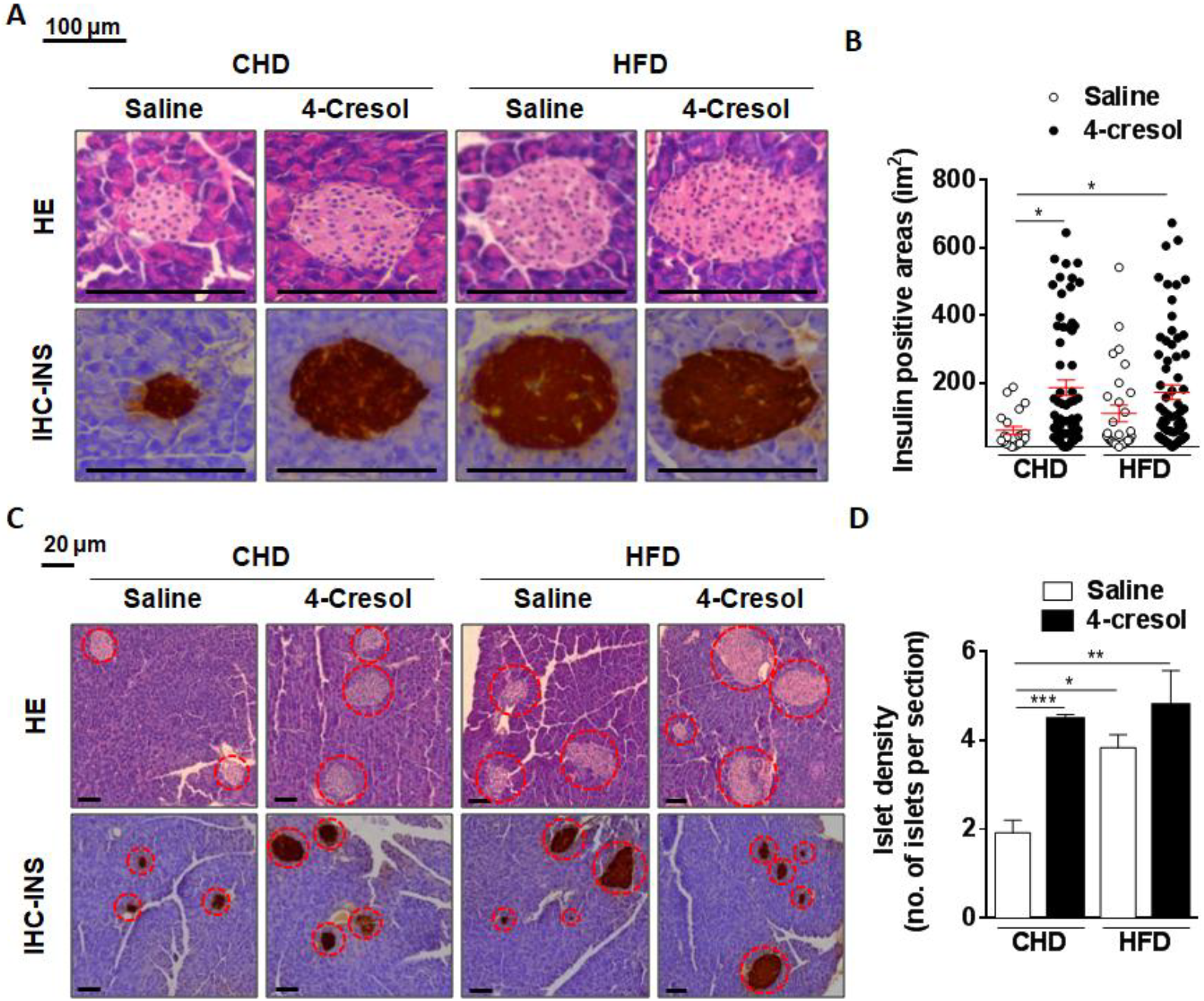
4-cresol treatment *in vivo* increases islet density and pancreatic vascularisation. Mice were fed normal chow diet (CHD) or high fat diet (HFD) and treated with chronic infusion of either 4-cresol or saline for 6-weeks. Pancreas sections were labeled either with Hematoxylin-Eosin (HE) and Immunohistochemistry (IHC) (A) to determine insulin positive area (B) and islet density (C,D). Results are means ± SEM. *P<0.05; **P<0.01; ***P<0.001; significantly different to relevant controls.

**Figure 5.**
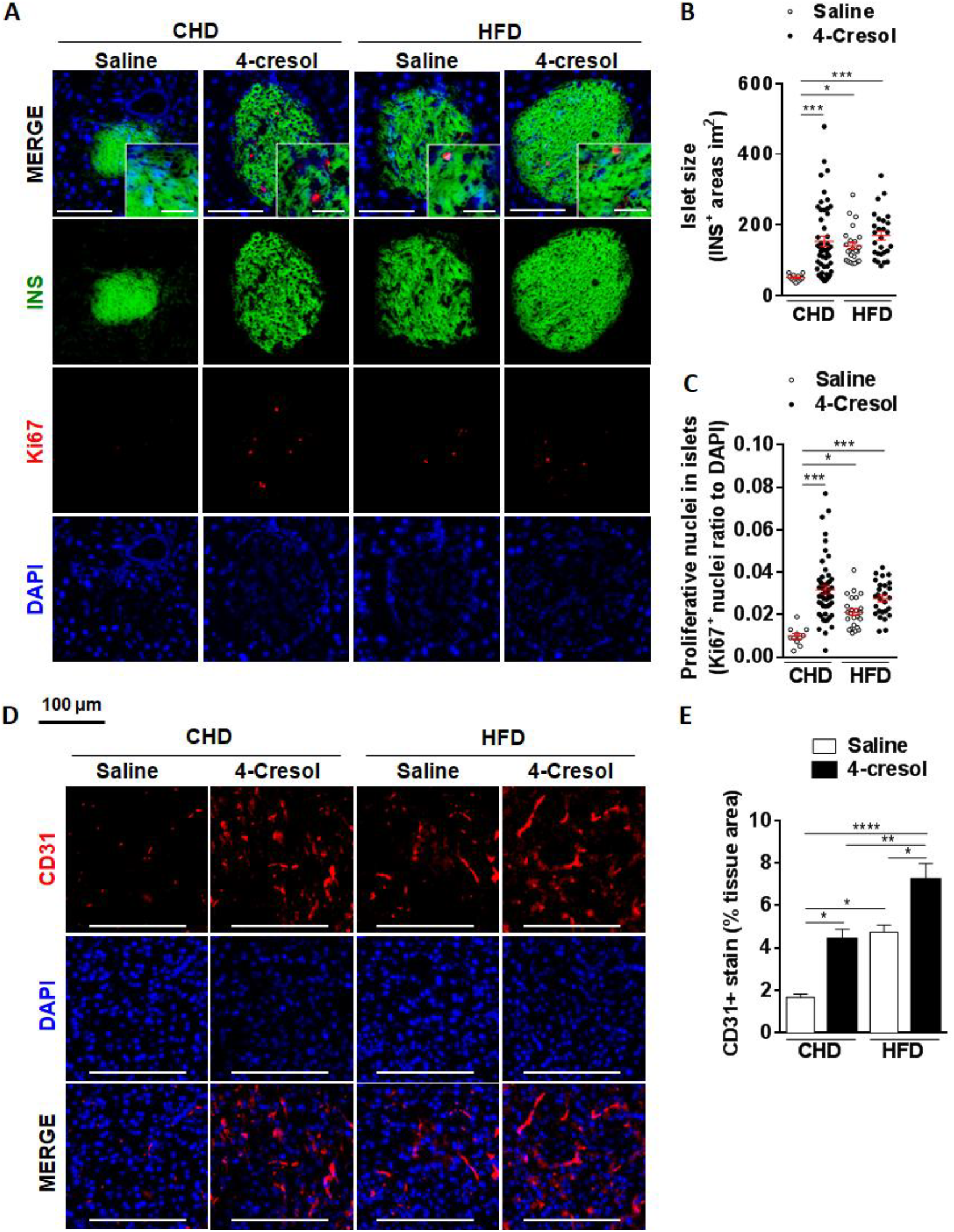
Chronic treatment of mice by 4-cresol promotes pancreas cell proliferation and vascularisation. Mice were fed normal chow diet (CHD) or high fat diet (HFD) and treated with chronic infusion of either 4-cresol or saline for 6-weeks. Pancreas sections were treated with Ki67 and DAPI to stain and quantify proliferative nuclei (A-C) and CD31 and DAPI to stain and quantify endothelial cells (D,E). Results are means ± SEM. *P<0.05; **P<0.01; ***P<0.001; ****P<0.0001 significantly different to relevant controls.

Collectively, these results illustrate the wide spectrum of pancreatic histological features and mechanisms affected by *in vivo* 4-cresol chronic administration, which may account for its effects on increased pancreas weight, enhanced glucose-stimulated insulin secretion *in vivo* and improved glucose tolerance in a model of obesity and insulin resistance induced experimentally.

### 4-cresol stimulates insulin content and cell proliferation in isolated mouse islets

To confirm the functional role of 4-cresol in pancreatic islets, we incubated islets isolated from mice with a concentration of 4-cresol (10nM) corresponding to the dose administered *in vivo* in mice, and a higher dose (100nM). The lower dose of 4-cresol induced a strong increase in insulin release under basal condition (2.8mM glucose) (+17.3%), a marked stimulation of insulin secretion in response to glucose 16.6mM (+25.8%, p=0.06) (**Figure 6A**), and a significant increase in islet insulin content (+33.7%, p<0.05) (**Figure 6B**). Incubation with 4-cresol at 100nM had no effect on insulin production, secretion and content. Labeling of islets with KI67 revealed a significant effect of 4-cresol 10nM on increased islet cell proliferation (**Figure 6C,D**), as illustrated in **Figure 6E**. These results demonstrate the role of 4-cresol on beta cell function *in vitro*, and corroborate *in vivo* data in mice treated chronically with 4-cresol.

**Figure 6.**
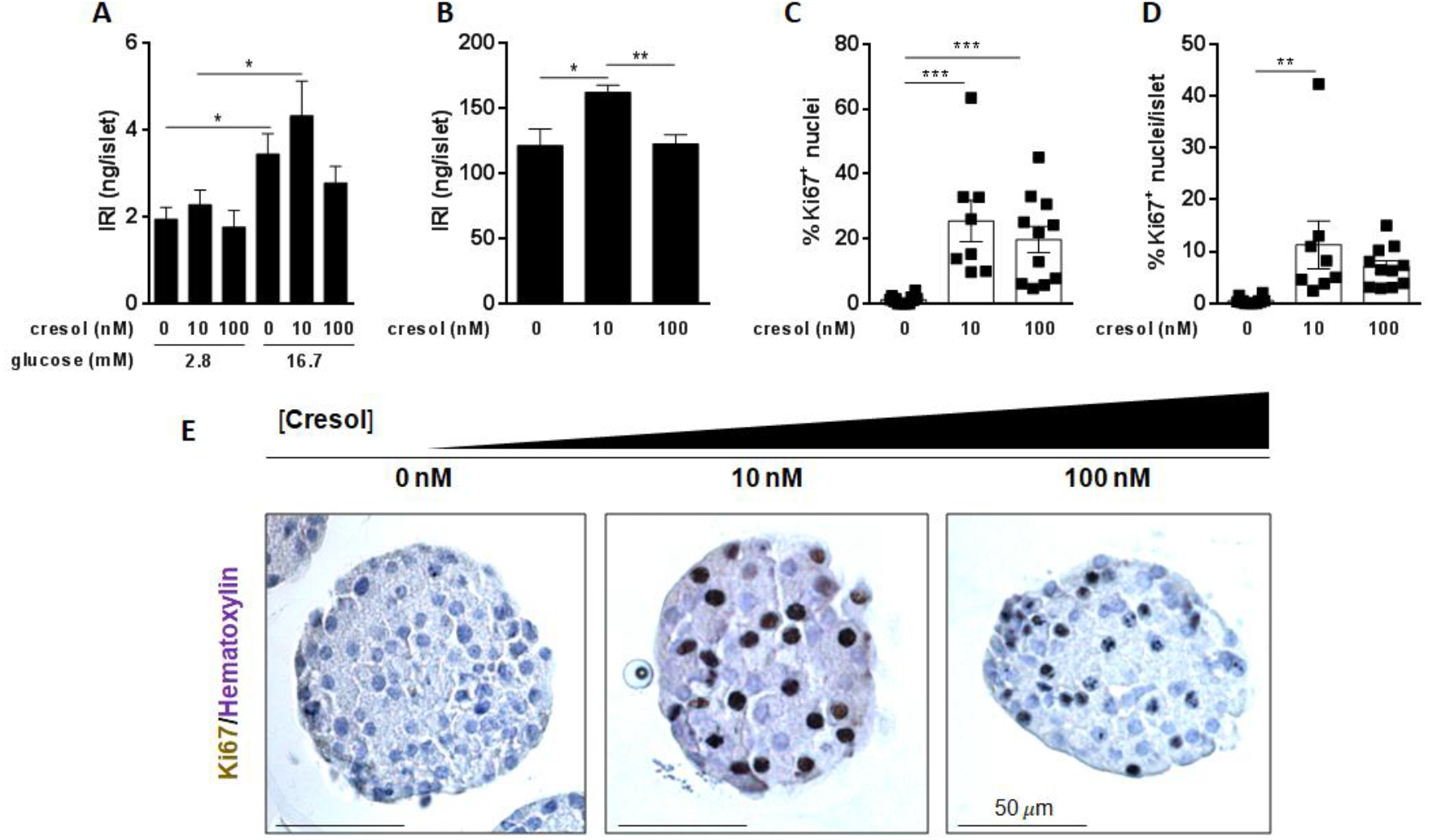
Incubation of islets with 4-cresol stimulates insulin content and cell proliferation. Insulin release and secretion in response to glucose (A) and insulin content (B) were determined in islets incubated with a control medium and with solutions of 4-cresol 10nM and 100nM. Sections of isolated islets treated with a medium free of 4-cresol and with 4-cresol 10nM and 100M were treated with Ki67 to quantify proliferative nuclei (C,D) illustrated in E. Results are means ± SEM. *P<0.05; **P<0.01; ***P<0.001 significantly different between groups.

### 4-cresol treatment improves glucose homeostasis and boosts insulin secretion and islet density in the Goto-Kakizaki rat

The impact of 4-cresol on enhanced insulin secretion and stimulated islet cell proliferation *in vivo* in a model of diet-induced insulin resistance and *in vitro* in isolated islets prompted us to test 4-cresol in a preclinical model of diabetes characterized by spontaneously-occurring reduction of β-cell mass. We used the Goto-Kakizaki (GK) rat, which exhibits glucose intolerance and deteriorated islet structure as a result of repeated breeding outbred Wistar rats over many generations using glucose intolerance for selecting breeders ^16^. Chronic administration of 4-cresol had no effect on body weight and BMI in this model of non obese diabetes (**Figure S4**). In contrast, adiposity index was significantly reduced, even though the GK is not a model of obesity, and pancreas weight nearly doubled (+94.6%, P<0.01) in 4-cresol treated rats (**Figure 7A,B**). This effect of 4-cresol was associated with a significant reduction in fasting glycemia (**Figure 7C**) and in glucose intolerance, assessed by a decrease in the glycemic response to the glucose challenge throughout the IPGTT (**Figure 7D**) and the significant drop in cumulative glycemia (**Figure 7E-F**). Fasting insulinemia and glucose-induced insulin secretion during the IPGTT were significantly more elevated in GK treated with 4-cresol than in rats treated with saline (**Figure 7G**). Pancreas histology analyses showed a significant increase in insulin positive area in response to 4-cresol, which was associated with increased cell proliferation determined by Ki67 labeling (**Figure 7H-J**). These results strongly support data obtained in HFD-fed mice and demonstrate that 4-cresol administration dramatically improves diabetes phenotypes in a model characterized by spontaneously occurring insulin deficiency and deteriorated islet structure.

**Figure 7.**
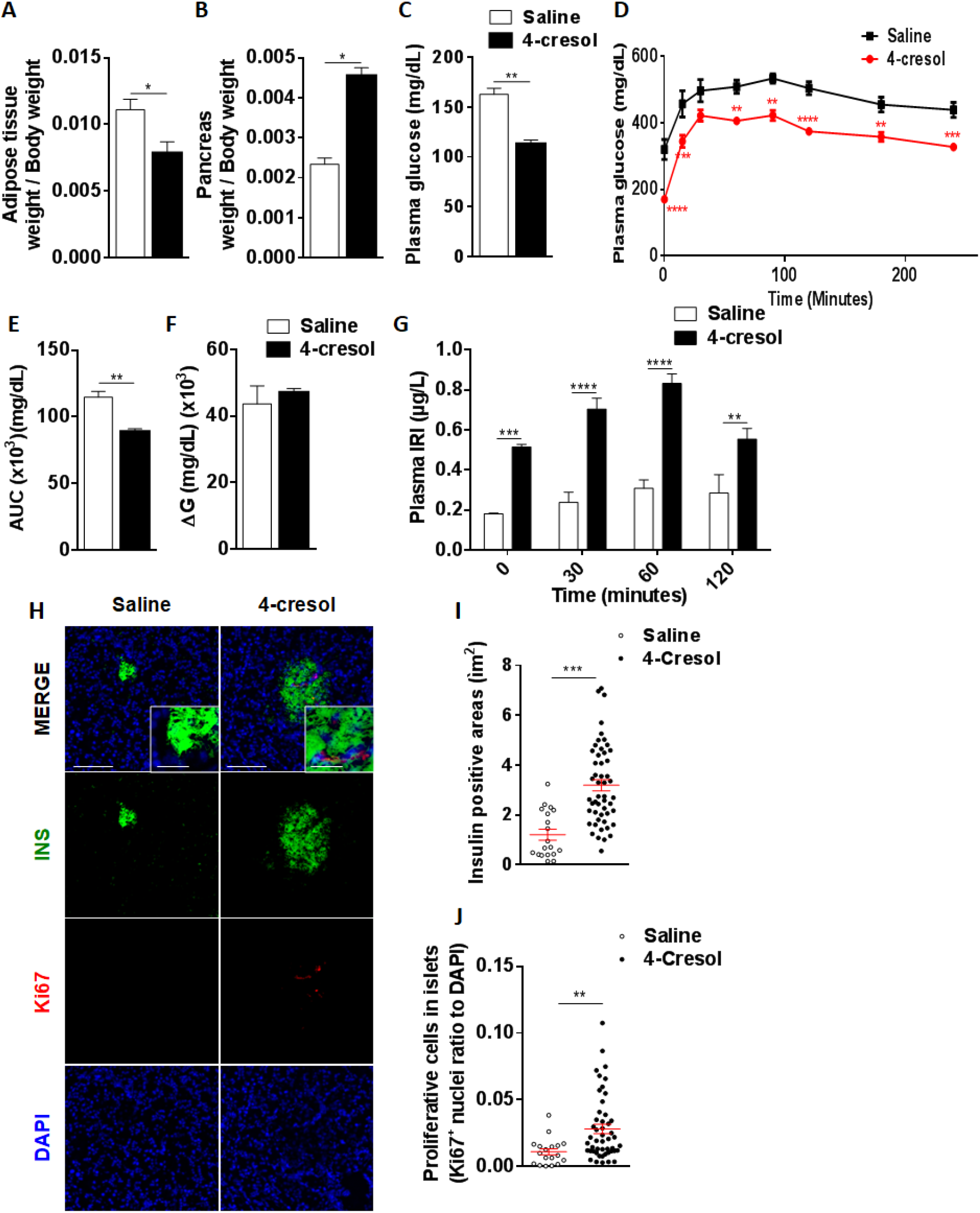
4-cresol treatment *in vivo* in Goto-Kakizaki rats improves glucose homeostasis and boosts insulin secretion and islet density. Adipose tissue and pancreas weight (A,B), fasting glycemia (C), glucose tolerance (D-F), glucose-stimulated insulin secretion (G) and pancreas histopathology (H-J) were determined in rats of the Goto-Kakizaki (GK) model of type 2 diabetes chronically treated with 4-cresol for 6-weeks. Pancreas sections were labeled either with Hematoxylin-Eosin and Immunohistochemistry to determine insulin positive area (I) and treated with Ki 67 and DAPI to stain and quantify proliferative nuclei (J). AUC was calculated as the sum of plasma glucose values during the IPGTT. ΔG is the AUC over the baseline value integrated over the 240 minutes of the test. All measures are from 6 rats per group. Data were analyzed using the unpaired Mann-Whitney test. Results are means ± SEM. **P<0.01; ***P<0.001; ****P<0.0001, significantly different to GK rats treated with saline. IRI, immunoreactive insulin.

### 4-cresol chronic treatment is associated with profound changes in pancreas gene expression

To get insights into molecular mechanisms that contribute to structural and functional changes induced by 4-cresol in the pancreas *in vivo*, expression of selected genes covering various aspects of pancreas biology was tested by quantitative RT-PCR in HFD-fed and control mice (**Figure 8A**) and in GK rats (**Figure 8B**). Expression of uncoupling protein 2 (*Ucp2*), interleukins (*Il6*, *Il10*) and tumor necrosis factor (*Tnfa*) was increased by HFD feeding. Chronic treatment by 4-cresol in CHD-fed mice led to increased transcription of genes encoding amylase, vascular endothelial growth factor (*Vegf*), brain derived neurotrophic factor (*Bdnf*), *Il10* and *Sirt1*, which coincided with increased NAD/NADH ratio, when compared to saline-treated CHD-fed mice. Enhanced expression of the insulin gene (*Ins1*) and the transcription factor HNF1 homeobox A (*Hnf1α*) by 4-cresol and reduced expression of *Ucp2* in this comparison were not statistically significant. In mice fed HFD 4-cresol induced significant overexpression of genes encoding Sirt1 and Ins1, and significantly reduced expression of *Ucp2*, *Il6* and *Tnfa* when compared to HFD-fed mice treated with saline. *Il10* transcript level remained elevated in HFD-fed mice treated with 4-cresol. *Bndf*, *Hnf1α* and *Vegf* were also strongly overexpressed in response to 4-cresol in these mice but differences to saline treated mice fed HFD were not statistically significant.

**Figure 8.**
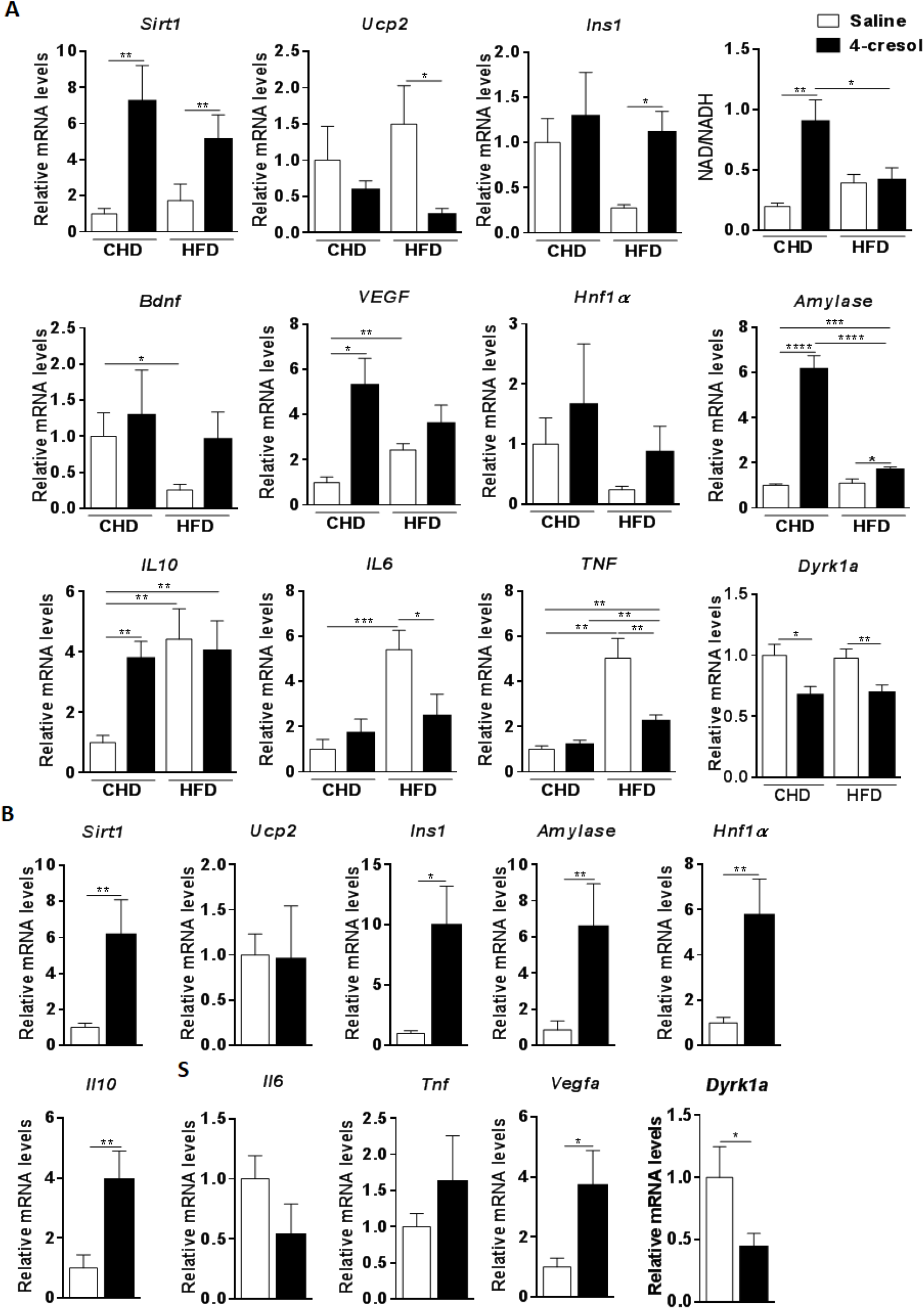
Effects of chronic administration of 4-cresol in fat fed mice and Goto-Kakizaki rats on pancreas gene expression. Transcription level of key genes covering various functions relevant to pancreas biology were determined by quantitative RT-PCR and the level of NAD and NADH were measured in the total pancreas of C57BL/6J mice fed control chow (CHD) or high fat diet (HFD) (A) and in Goto-Kakizaki (GK) rats (B). Results are means ± SEM. *P<0.05; **P<0.01; ***P<0.001; ****P<0.0001, significantly different to relevant controls.

The stimulatory effects of 4-cresol chronic administration on the expression of *Sirt1*, *Ins1*, *Hnf1α*, *Il10* and *Vegfa* were replicated in GK pancreas (**Figure 8B**). Expression of genes encoding amylase, *Il6* and *Tnfa* was strongly altered in 4-cresol treated GK rats, but differences to controls treated with saline were not statistically significant.

Due to strong similarities in the biological effects of 4-cresol and harmine, an inhibitor of the dual specificity tyrosine phosphorylation regulated kinase (DYRK1A) known to stimulate β-cell proliferation ^17,18^, we tested the expression of this gene in fat fed mice and GK rats treated with 4-cresol. In both models, 4-cresol consistently downregulated pancreatic expression of DYRK1A, thus suggesting that 4-cresol impacts signaling pathways similar to those mediating the cellular action of other natural products, such as harmine.

These results demonstrate the broad ranging molecular consequences of 4-cresol treatment in the pancreas of animal models of diabetes induced experimentally by dietary changes or caused by naturally occurring genetic polymorphisms, and provide insights into possible mediating mechanisms.

## DISCUSSION

We report correlations between type 2 diabetes (T2D) and obesity and variations in serum concentration of a series of metabolites derived by GC-MS based metabolomics. Significant associations were found with metabolic products of gut microbial metabolism, including 4-cresol, which was tested for its biological role *in vivo* in preclinical models of spontaneous or experimentally-induced diabetes and *in vitro* in β-cells. We demonstrate that 4-cresol improves glycemic control, enhances glucose-stimulated insulin secretion, reduces adiposity and liver triglycerides and stimulates pancreatic islet density, vascularisation and cell proliferation *in vivo* and *in vitro*. These effects mimic metabolic improvements induced by harmine, and may be mediated through downregulated expression of DYRK1A.

Our metabolomic approach identified systematic correlations between T2D and obesity and a metabotype ^7^ of 18 co-associated metabolites, which covers obvious candidates for these conditions, including for example glucose, branched chain amino acids and mannose, which was recently associated with obesity and is proposed as an accurate marker of insulin resistance ^19^. Glucose-and mannose-mediated inhibition of cellular uptake of 1,5-anhydro-D-sorbitol ^20^, a marker of short-term glycemic control ^21^, supports the strongly significant negative correlation between serum 1,5-anhydro-D-sorbitol and T2D and obesity in our study. The metabotype significantly correlated with these diseases also includes 2-hydroxybutyric acid, which is elevated in plasma of T2D patients ^22^ and could be a predictive marker of insulin resistance ^23^, and 2-amino adipic acid which is associated with T2D risk ^24^. Stimulation of insulin secretion *in vitro* ^24^ by 2-amino adipic acid and its regulation by the dehydrogenase E1 and transketolase domain-containing 1 gene (DHTKD1), which plays critical roles in diabetes pathogenesis ^25^ and in mitochondrial energy production and biogenesis ^26^, suggest broad ranging roles of 2-amino adipic acid in the pathogenesis of metabolic disorders.

There is growing interest in understanding the impact of structural changes in gut bacterial communities on host metabolism ^1,11^ and disease pathogenesis ^2,27^. We demonstrate significant correlations between the microbial metabolite 4-cresol and T2D and obesity. 4-cresol is a product of colonic fermentation of tyrosine and phenylalanine ^28^, which can be also synthesised from phenol found in the environment and absorbed by ingestion, inhalation and dermal contact. It is present in food (smoked foods, tomatoes, asparagus, dairy products), drinks (coffee, black tea, wine), cigarette smoke, wood burning and surface waters and groundwater (ATSDR, www.atsdr.cdc.gov/). Lethal dose (LD50) for 4-cresol given orally ranges from 200 to 5000 mg/kg/day ^13^. Irritation to the respiratory epithelium and deteriorated liver function can be caused by dietary exposure to doses of 4-cresol (240-2000mg/kg/day) which are much higher than those used in our *in vivo* (0.5mg/kg/day) and *in vitro* (10nM) studies.

Results from chronic administration of low dose of 4-cresol *in vivo* in fat diet fed mice and Goto-Kakizaki rats provided evidence for its broad ranging biological roles and its beneficial effects on glucose homeostasis and β-cell function. They did validate the inverse correlation of this metabolite with T2D and obesity in humans in our study. Consistent with our *in vivo* findings, 4-cresol was reported to inhibit proliferation and differentiation of 3T3-L1 preadipocytes ^29^. Also, negative association between BMI and urine concentration of its detoxification product 4-cresyl sulfate in humans ^30^ suggests physiological roles of other elements in the cresol pathway. Conserved effects of 4-cresol and 4-MC, one of its potential products through dehydrogenation by bacterial metabolism ^14^, on glucose tolerance, insulin secretion, adiposity and body and organ weights support this observation. This is further supported by the reduction of β-cell apoptosis and hyperglycemia mediated by 4-MC in diabetic rats ^31^. These results suggest low specificity of 4-cresol and 4-MC which may regulate similar signalling systems.

Our findings underline the broad ranging impact of 4-cresol on improved glucose regulation, enhanced insulinemia, glucose-induced insulin secretion *in vivo* and insulin content *in vitro*, reduced liver triglycerides and possibly islet neogenesis. In addition, results from pancreas histopathology and gene expression analysis in both preclinical models suggest that 4-cresol contributes to reduce inflammation, as illustrated by both downregulated expression of *Il6* and *Tnfa* and stimulated expression of *Il10*, an anti-inflammatory marker involved in vascularisation and resistance to apoptosis ^32,33^. Upregulated expression of *Vegf* and *Bdnf* suggests enhanced vascularisation and parasympathetic tone, respectively. Of note, *Bdnf* reduces hyperglycemia and increases the number and area of pancreatic islets in diabetic *db/db* mice ^34^.

*Sirt1* upregulated pancreatic expression by 4-cresol in mice and GK rats, alongwith increased NAD^+^/NADH ratio and downregulated expression of *Ucp2* in mice, suggest that it may mediate the effects of 4-cresol. SIRT1 stimulates insulin secretion and lipolysis, reduces adipogenesis, liver lipid accumulation and inflammation, and improves glucose homeostasis ^35^. SIRT1 is regulated by the polyphenol resveratrol, which reduces adiposity through stimulation of lipolysis and inhibition of lipogenesis, exhibits anti-inflammatory effects through downregulated expression of *Tnfa* and *Il6* ^36^, and potentiates insulin secretion in insulinoma cells and human islets ^37^.

These data suggest convergent biological roles of natural products or polyphenols and the phenol derivative 4-cresol, which may share ligands and signalling mechanisms leading to improved glucose homeostasis. We showed that 4-cresol and harmine exhibit very similar effects on increased islet mass and improved glucose regulation ^38^. Harmine directly targets the kinase DYRK1A ^39^ resulting in its inhibition and in increased human β-cell proliferation, β-cell mass and insulin content ^17,18^. Interestingly, the role of polyphenols on DYRK1a inhibition is suggested by the rescue of neurobehavioral deficits in transgenic mice overexpressing DYRK1a through treatment with green tea polyphenols ^40^. Our results indicate that 4-cresol is also a DYRK1A inhibitor and support the emerging role of kinases in mediating the cellular function of microbial metabolites ^41^.

In conclusion, our findings illustrate the power of systematic metabolic profiling to identify a metabotype integrating microbial metabolites that are biological endpoints of gut microbiome architecture and reflect gut microbiota function in chronic diseases. We demonstrate the beneficial role of low doses of 4-cresol on β-cell function and diabetes phenotypes. Deeper exploitation of the metabolomic dataset beyond targeted analysis of known molecules may uncover networks of metabolites that are co-regulated with 4-cresol and can also improve diabetes and obesity variables, and point to underlying regulatory gene pathways. The identification of bacterial ecosystems that synthesize 4-cresol and its co-metabolites may have important therapeutic relevance in the treatment of T2D. Our results pave the way to the identification of diagnostic and prognostic metabolic biomarkers at the crossroads between diabetes risk and microbiome activity, as well as information of genes, biological pathways and bacterial species contributing to 4-cresol metabolism that can provide therapeutic solutions in diabetes.

## Acknowledgements

The authors acknowledge financial support from the European Commission for collection of the patient cohort (FGENTCARD, LSHGCT-2006-037683) and experimental work in mice (METACARDIS, HEALTH-F4-2012-305312).

## Author Contributions

PZ, FM, JKN, ML, MED and DG conceived the study. PZ provided patients serum samples. KS, TAS and FM carried out targeted GC-MS metabolomic analyses. FB, ALL and NP carried out mouse experiments. FB, KM and CM performed incubation of isolated islets. FB, ALL and FA performed histology and gene expression analyses. JC analysed pharmacological data. LH and ARM performed statistical analyses. PZ and DG wrote the manuscript. All authors have given approval to the final version of the manuscript.

## Declaration of Interests

Conflicts of interest: FB, FM, PZ and DG are named inventors on a patent related to this work (Ref. EP 17306326). KS and TAS are employees of Shimadzu, Kyoto, Japan. FA, KM, ALL, NP, JC, LH, ARM, NV, JKN, CM, ML and MED declare no competing financial interests.

## Materials and Methods

Full details of Material s and Methods are available in Supplementary Material

### Study Subjects

We used plasma samples from 138 subjects of the FGENTCARD collection ^42^.

### Gas chromatography coupled to mass spectrometry (GC-MS)

GC-MS analysis was performed using a GCMS-QP2010 Ultra (Shimadzu, Kyoto, Japan). By comparing mass spectral pattern of chromatographic peaks to those in the NIST library or Shimadzu Database, 101 metabolites were identified (Supplementary Table 1).

### Animal experiments

C57BL/6J mice were from a commercial supplier (Janvier Labs, Courtaboeuf, France). Goto-Kakizaki rats were bred locally. Mice were fed either control carbohydrate diet (D 12450Ki, Research diets, NJ) or high fat diet (HFD) (D12492i, Research diets, NJ) for a week until osmotic minipumps Alzet^®^ (Charles River Lab France, l’Arbresle, France) filled with saline, 4-cresol or 4-methylcatechol (Sigma Aldrich, St Quentin, France) were inserted subcutaneously (flow rate 0.15μL/h). The same procedure was applied in GK rats fed control diet. Intraperitoneal glucose tolerance tests were performed in overnight fasted animals. After six weeks of metabolite treatment, animals were killed and organs were either snap frozen or processed for histopathology. Procedures were carried out under national licence condition (Ref 00486.02).

### *In vitro* insulin secretion in mouse isolated islets

Islets were isolated in a solution containing collagenase (Roche, Sigma Aldrich, St Quentin, France) and purified on a four layer density gradient of Histopaque 1119 (Sigma-Aldrich, France). Islets incubated with the culture medium or cresol (10nM or 100nM) (W233706, Sigma Aldrich, St Quentin, France) were either prepared to determine insulin content or incubated with 2.8 or 16.7mM glucose.

### Histology of animal tissues and isolated islets

Tissue sections were stained in Hematoxylin and Eosin. Antibodies were used for immunohistochemistry detection of insulin on pancreas sections (Dako, Saint Aubin, France). Liver sections were incubated with an Oil red O staining solution (Sigma Aldrich, St Quentin, France). For double immunostaining and immunofluorescence, pancreas sections were co-stained with primary Anti-Ki67 antibody and secondary Donkey Anti-Goat IgG H&L conjugated to Alexa Fluor^®^ 568, or with primary Mouse/Rat CD31/PECAM-1 Antibody and secondary Goat anti-Rabbit IgG H&L conjugated to Alexa Fluor 488. For islet immunohistochemistry, sections prepared in agarose containing heamatoxylin were incubated with primary antibodies against Ki67 (Abcam, Paris, France) and with HRP-conjugated secondary antibody (Bio-Rad, Marnes-la-Coquette, France).

### Gene expression

Quantitative RT-PCR was performed using SYBR green assays (Eurogentec, Angers, France). Primers are listed in Supplementary Table 2.

### DYRK1A analyses

4-cresol binding to DYRK1A was assessed as previously described ^43,44^. The dissociation constant (Kd) was calculated from duplicate 12-point dose–response curve. The DYRK1A half-maximal inhibitory curve (IC50) for 4-cresol was determined using Kinexus service (Kinexus, Vancouver, Canada).

### Analytical assays

Blood glucose was measured using Accu-Check^®^ Performa (Roche Diagnostics, Meylan, France) and plasma insulin was determined by ELISA (Mercodia, Uppsala, Sweden). Liver triglycerides were determined using a colorimetric assay (Abcam, Paris, France). NAD/NADH was determined using a quantification kit (Sigma Aldrich, St Quentin, France).

### Statistical analyses

R statistics was used to compute P-values for correlations between metabolic features and clinical data. Multivariate modelling of the signature related to diabetes and obesity was performed through Orthogonal Partial Least Squares Discriminant Analysis ^45^. All p-values were corrected for multiple testing using the Benjamini-Hochberg procedure.

---

**Figure S1.**
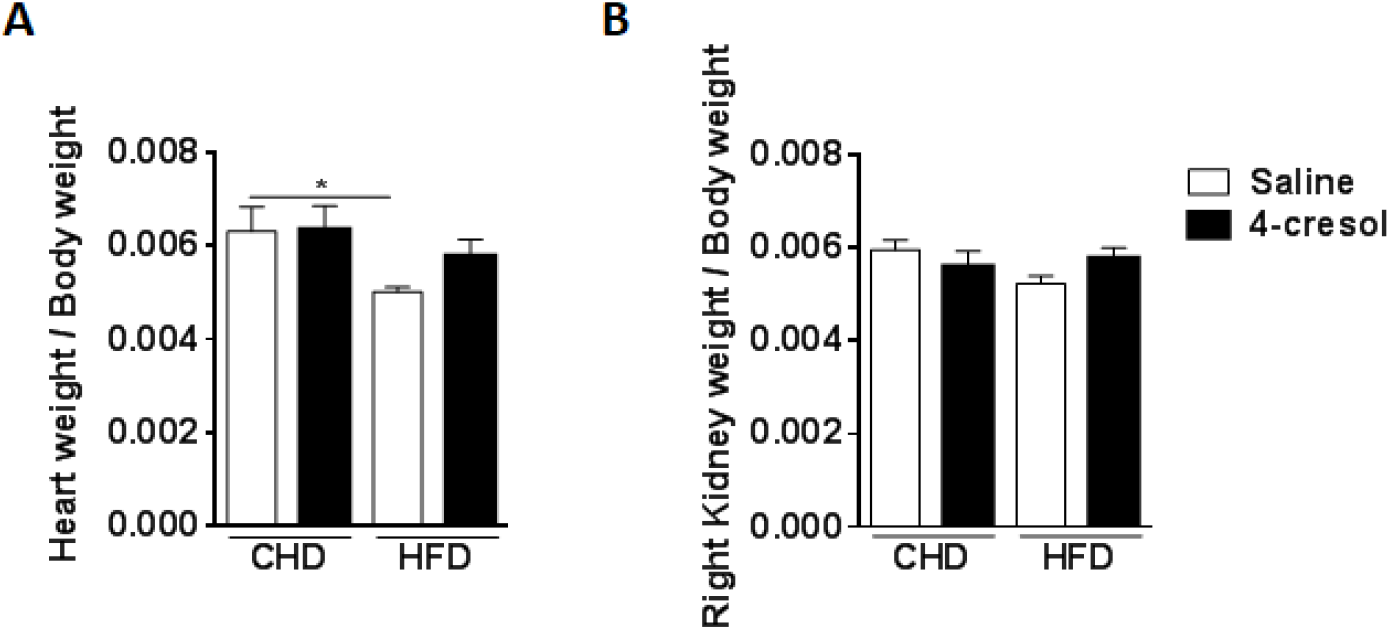
Effects chronic treatment of C57BL6/J mice *in vivo* 4-cresol on heart weight (a) and kidney weight (B). Data were analysed using the unpaired Mann-Whitney test. Results are means ± SEM. *P < 0.05 significantly different to relevant controls.

**Figure S2.**
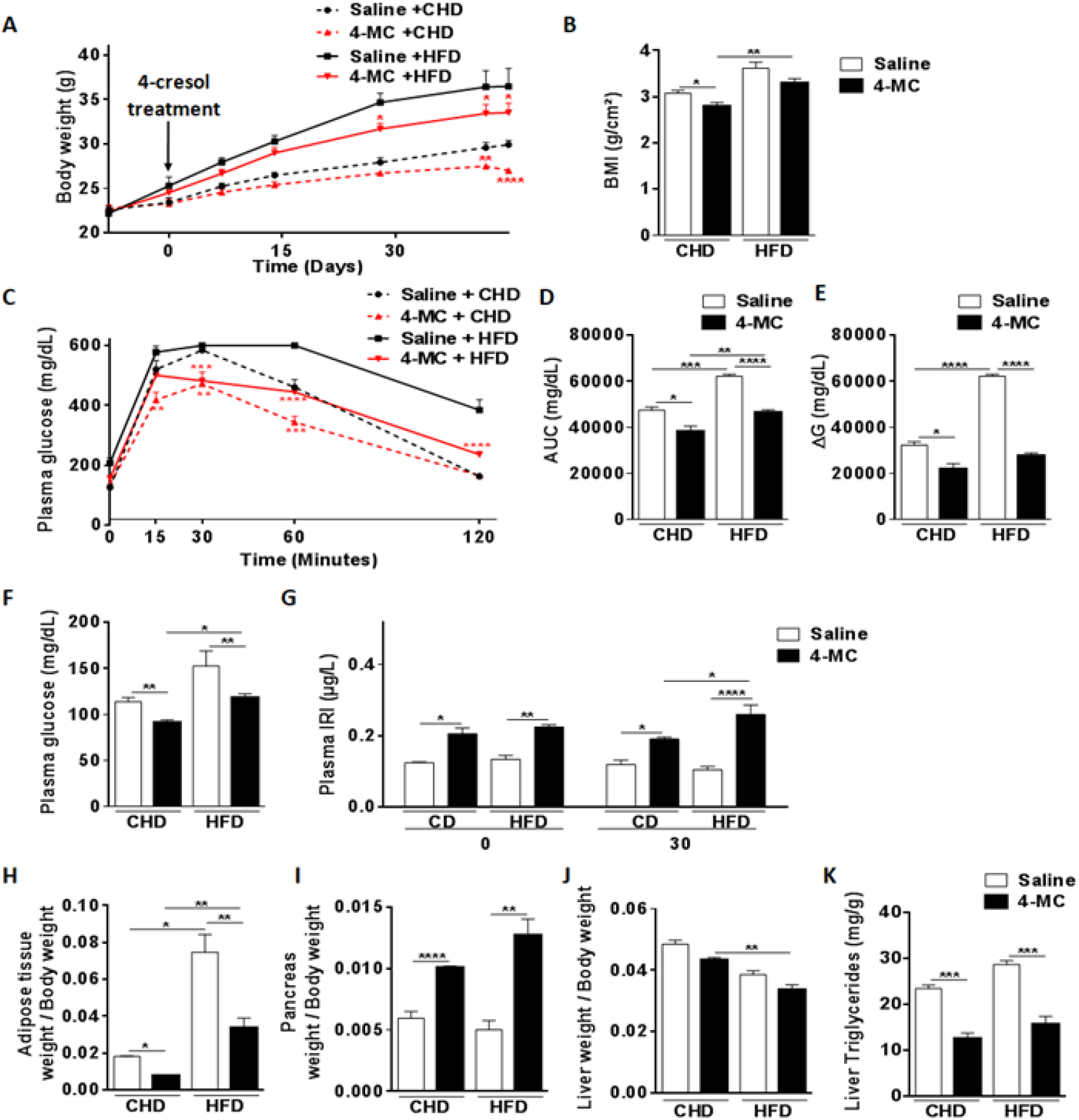
4-cresol derivative 4-methylcatechol mimics the effects of 4-cresol on glucose homeostasis, insulin secretion and obesity in C57BL6/J mice. The effects of 6-week 4-methylcatechol (4-MC) treatment *in vivo* in mice fed control chow or high fat diet (HFD) were tested on body weight (A), body mass index (BMI) (B), glucose homeostasis (C-F), glucose-stimulated insulin secretion (G), organ weight (H-J) and liver triglycerides (K). BMI was calculated as body weight divided by the squared of anal-nasal length. AUC was calculated as the sum of plasma glucose values during the IPGTT. ΔG is the AUC over the baseline value integrated over the 120 minutes of the test. All measures are from 6 mice per group. Data were analysed using the unpaired Mann-Whitney test. Results are means ± SEM. *P < 0.05; **P < 0.01; ***P < 0.001;****P < 0.0001, significantly different to relevant controls.

**Figure S3.**
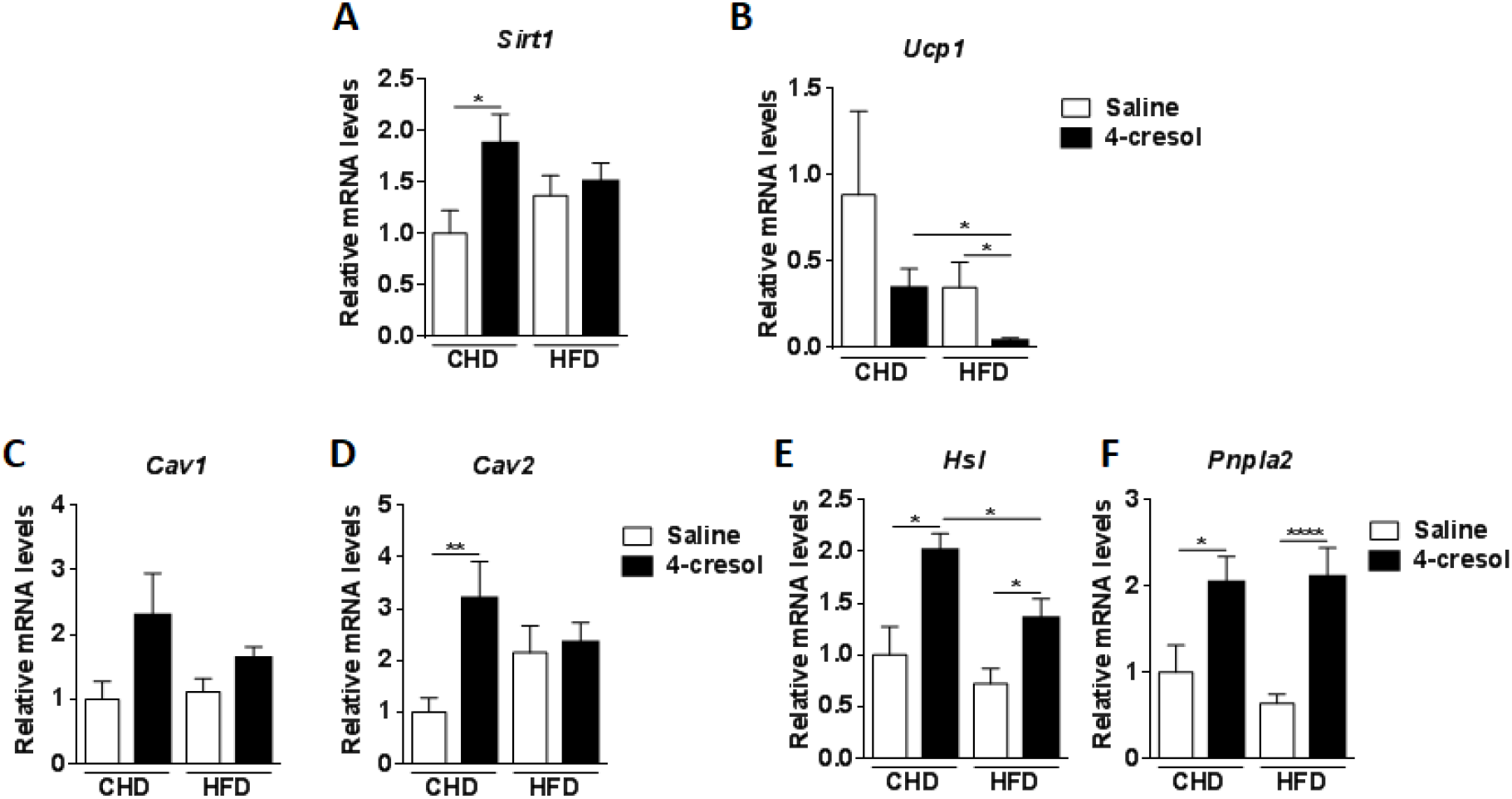
Effects of chronic treatment of mice by 4-cresol *in vivo* on gene expression in adipose tissue. Transcript abundance of key genes regulating adipose tissue biology was tested by quantitative RT-PCR in retroperitoneal fat pads of mice fed control chow diet (CHD) or high fat diet (HFD). Data were analysed using the unpaired Mann-Whitney test. Results are means ± SEM. *P < 0.05; **P < 0.01; ****P < 0.0001, significantly different to relevant controls.

**Figure S4.**
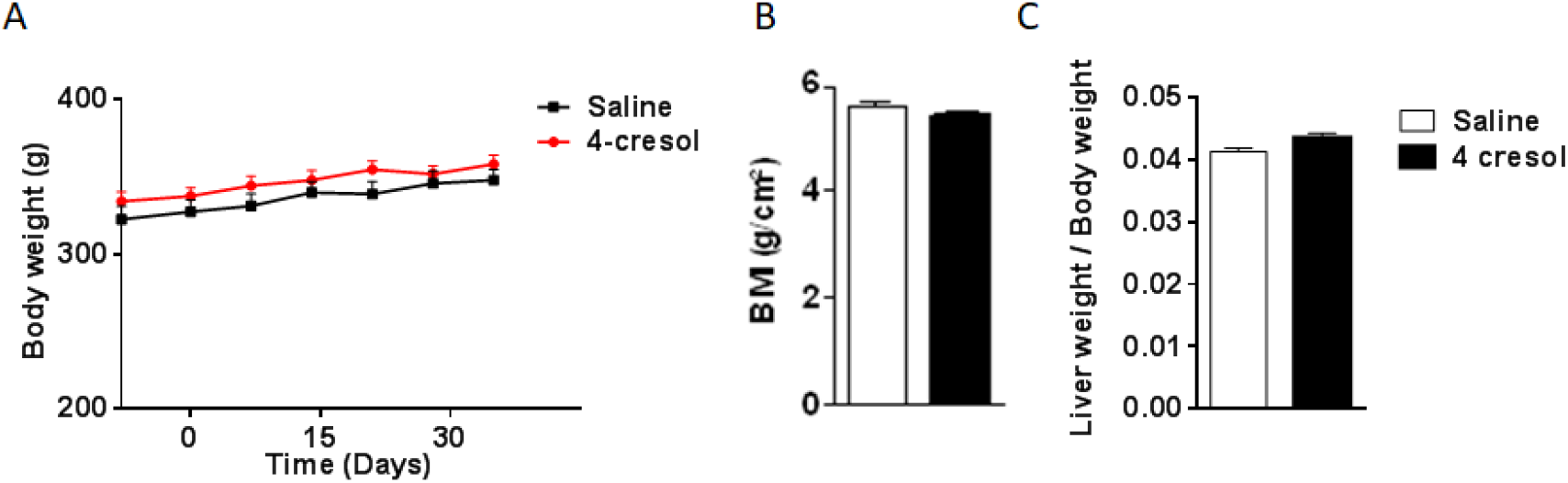
Effects of chronic administration of 4-cresol in vivo in Goto-kakizaki (GK) rats on body weight and organ weight. The effects of 6-week 4-cresol treatment *in vivo* in GK rats were tested on body weight(A), body mass index (BMI) (B) and liver weight to body weight ratio (C). BMI was calculated as body weight divided by the squared of anal-nasal length. All measures are from 6 rats per group. Data were analysed using the unpaired Mann-Whitney test. Results are means ± SEM.

**Supplementary Table 1.** Mass spectrometry features of metabolites analysed in serum samples in the cohort of coronary artery disease patients (n=77) and controls (n=61). A typical spectrum is shown in Supplementary Figure 1.

|  | Quantitative ion | Qualifier ion | Retention time (min) |
| --- | --- | --- | --- |
| Pyruvic acid | 174 | 115 | 7.495 |
| Lactic acid | 219 | 101 | 7.653 |
| Phenol | 166 | 151 | 7.712 |
| Glycolic acid | 205 | 177 | 7.796 |
| 2-Oxobutyric acid | 188 | 89 | 8.118 |
| Alanine | 218 | 190 | 8.168 |
| 3-Methyl-2-oxobutyric acid | 89 | 100 | 8.223 |
| Oxalic acid | 219 | 190 | 8.39 |
| 2-Hydroxybutyric acid | 131 | 205 | 8.434 |
| 3-Hydroxybutyric acid | 191 | 117 | 8.778 |
| 3-Hydroxyisobutyric acid | 177 | 218 | 8.778 |
| 4-cresol (p-Cresol) | 180 | 165 | 8.822 |
| 2-Hydroxyisovaleric acid | 145 | 219 | 8.878 |
| 2-Aminobutyric acid | 130 | 218 | 8.955 |
| 3-Hydroxyisovaleric acid | 131 | 205 | 9.349 |
| 2-Oxoisocaproic acid | 200 | 216 | 9.368 |
| Valine | 218 | 100 | 9.452 |
| Urea | 204 | 171 | 9.554 |
| Octanoic acid | 201 | 129 | 9.884 |
| Glycerol | 218 | 205 | 9.956 |
| Leucine | 232 | 218 | 10.007 |
| Phosphoric acid | 225 | 283 | 10.028 |
| 2-Aminoethanol | 174 | 262 | 10.035 |
| Acetylglycine | 145 | 130 | 10.069 |
| Isoleucine | 232 | 218 | 10.254 |
| Succinic acid | 247 | 172 | 10.363 |
| Proline | 216 | 244 | 10.382 |
| Glycine | 174 | 248 | 10.468 |
| Glyceric acid | 292 | 189 | 10.545 |
| Fumaric acid | 245 | 143 | 10.64 |
| Serine | 306 | 278 | 10.851 |
| Threonine | 291 | 218 | 11.129 |
| Glutaric acid | 261 | 158 | 11.205 |
| 2-Deoxytetronic acid | 321 | 233 | 11.47 |
| beta-Alanine | 290 | 174 | 11.579 |
| Decanoic acid | 229 | 117 | 11.74 |
| 3-Aminoisobutyric acid | 304 | 248 | 11.944 |
| Malic acid | 233 | 133 | 12.005 |
| Thereitol | 217 | 103 | 12.114 |
| Erythritol | 217 | 307 | 12.196 |
| Aspartic acid | 232 | 218 | 12.307 |
| Methionine | 250 | 293 | 12.416 |
| 4-Hydroxyproline | 140 | 230 | 12.424 |
| 5-Oxoproline | 258 | 230 | 12.483 |
| Threonic acid | 292 | 220 | 12.65 |
| Cysteine | 218 | 220 | 12.709 |
| 2-Oxoglutaric acid | 198 | 288 | 12.748 |
| Creatinine | 115 | 143 | 12.857 |
| IS(2-Isopropylmalic acid) | 275 | 349 | 12.938 |
| Glutamic acid | 246 | 363 | 13.125 |
| Lauric acid | 257 | 117 | 13.389 |
| Phenylalanine | 192 | 218 | 13.397 |
| Xylose | 307 | 217 | 13.484 |
| Arabinose | 307 | 217 | 13.533 |
| Asparagine | 231 | 188 | 13.596 |
| Pyrophosphate | 451 | 466 | 13.599 |
| Ribulose | 263 | 205 | 13.658 |
| Ribose | 307 | 217 | 13.66 |
| 2-Aminoethanesulfonic acid | 326 | 188 | 13.778 |
| 2-Aminoadipic acid | 260 | 128 | 13.894 |
| Xylitol | 217 | 103 | 13.937 |
| Arabitol | 217 | 103 | 14.027 |
| Aconitic acid | 375 | 285 | 14.083 |
| Indoxyl sulfate | 277 | 163 | 14.114 |
| Fucose | 117 | 160 | 14.164 |
| Phosphoglycerol | 445 | 357 | 14.279 |
| Glutamine | 362 | 347 | 14.386 |
| O-Phosphoethanolamine | 299 | 174 | 14.564 |
| Isocitric acid | 245 | 319 | 14.693 |
| Citric acid | 363 | 347 | 14.725 |
| Citrulline | 256 | 157 | 14.761 |
| Hypoxanthine | 265 | 280 | 14.78 |
| Ornithine | 420 | 174 | 14.799 |
| Myristic acid | 285 | 129 | 14.876 |
| Hippuric acid | 236 | 206 | 15.032 |
| 1,5-Anhydro-D-sorbitol | 259 | 362 | 15.053 |
| Fructose | 307 | 217 | 15.202 |
| Mannose | 319 | 160 | 15.33 |
| Lysine | 174 | 156 | 15.515 |
| Histidine | 371 | 356 | 15.565 |
| Glucose | 229 | 307 | 15.605 |
| Tyrosine | 382 | 354 | 15.668 |
| Glucuronic acid | 423 | 333 | 15.753 |
| Indol-3-acetic acid | 202 | 319 | 15.962 |
| Paraxanthine | 237 | 252 | 16.055 |
| Palmitoleic acid | 311 | 145 | 16.138 |
| Xanthine | 353 | 368 | 16.206 |
| Gluconic acid | 292 | 333 | 16.224 |
| Palmitic acid | 313 | 145 | 16.236 |
| scyllo-Inositol | 318 | 305 | 16.395 |
| Uric acid | 367 | 353 | 16.688 |
| 3-Indolepropionic acid | 202 | 333 | 16.787 |
| myo-Inositol | 432 | 318 | 16.826 |
| Margaric acid | 117 | 327 | 16.87 |
| Kynurenine | 307 | 424 | 17.361 |
| Indolelactic acid | 202 | 421 | 17.37 |
| Oleic acid | 339 | 129 | 17.374 |
| Tryptophan | 291 | 218 | 17.633 |
| Cystine | 411 | 266 | 17.957 |
| Sucrose | 361 | 217 | 20.174 |
| Maltose | 204 | 361 | 21.103 |

**Supplementary Table 2.**
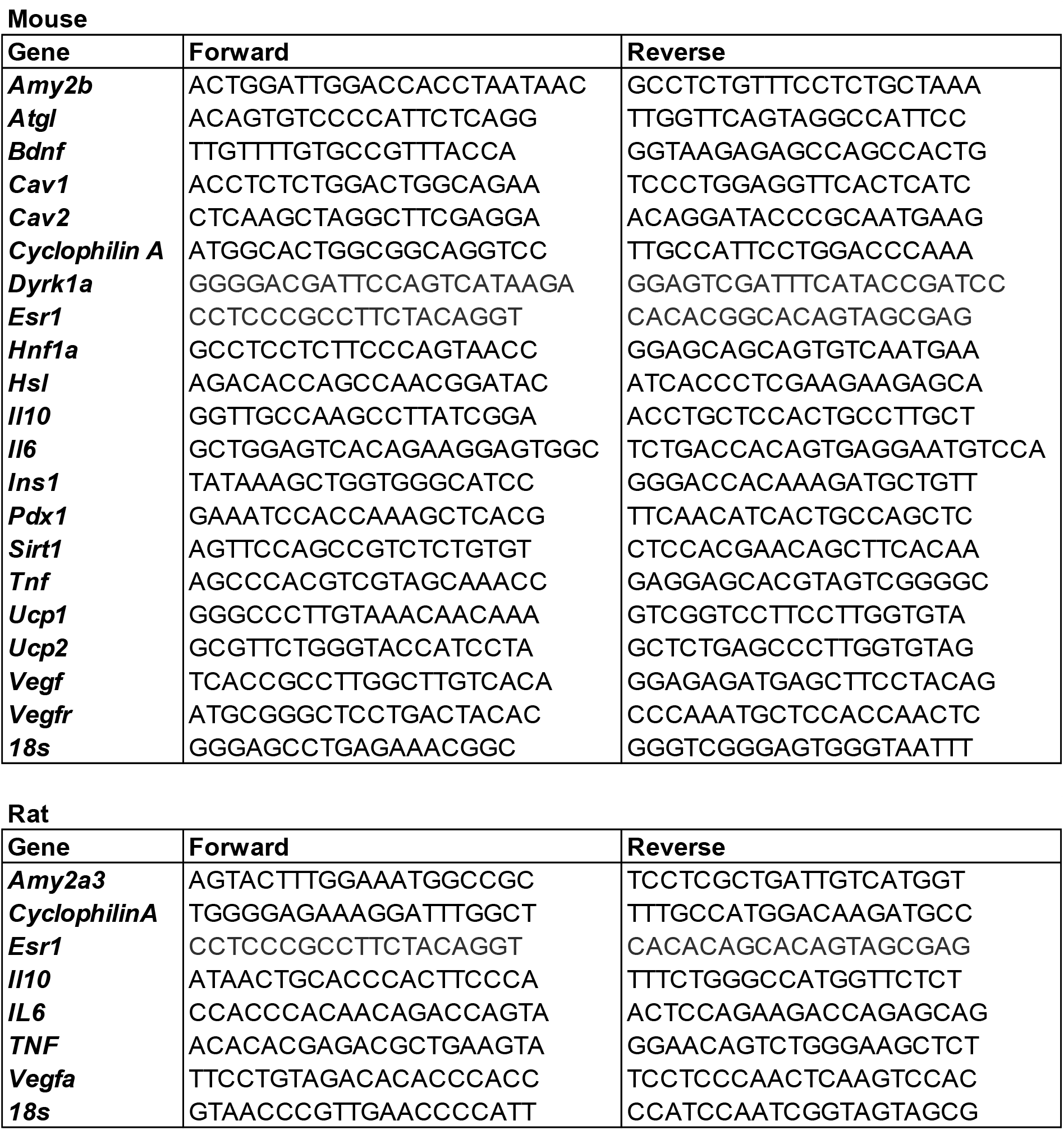
Oligonucleotides used in qPCR experiments in mice and rats

## Supplementary Methods

### Study Subjects

A total of 138 subjects (90 males and 48 females) recruited between 2006 and 2009 for inclusion in the FGENTCARD patient collection (*1*) were selected on the basis of extreme levels of plasma HDL cholesterol, presence of coronary stenosis and a positive family history of the disease, defined as a sibling, parent, or second-degree relative with a coronary event. Presence of diabetes, hypertension or obesity was recorded. These subjects were originally referred to a catheterization care unit for clinical evaluation. Patients provided a written consent for the whole study including genomic analyses. The study protocol was approved by the Institutional Review Board at the Lebanese American University.

### Chemicals

Purified water (TOC grade), methanol (residual PCB grade), chloroform (residual PCB grade), and pyridine dehydrated were purchased from Wako Pure Chemical Industries (Osaka, Japan). Methoxyamine hydrochloride and 2-isopropylmalic acid were obtained from Sigma-Aldrich (Sigma-Aldrich, Tokyo, Japan).

### Sample preparation

The internal standard solution (2-isopropylmalic acid, 0.1 mg/mL in purified water) and extraction solvent (methanol: water: chloroform = 2.5:1:1) were mixed at a ratio of 6:250, which was added to 50μL of each plasma sample. The resulting solution was mixed using a shaker at 1,200 rpm for 30 min at 37°C. After centrifugation at 16,000 x g for 5 min at 4°C, 150μL of supernatant was collected and mixed with 140μL of purified water. The solution was thoroughly mixed and centrifuged at 16,000 x g for 5 min at 4°C. Finally, 180μL of supernatant was collected and lyophilized. The lyophilized sample was dissolved in 80μL of methoxyamine solution (20 mg/mL in pyridine) and agitated at 1,200 rpm for 30 min at 37°C. Forty micro litters of N-methyl-N-trimethylsilyltrifluoroacetamide solution (GL science, Tokyo, Japan) were added for trimethylsilyl derivatization, followed by agitation at 1,200 rpm for 30 min at 37°C. After centrifugation at 16,000 x g for 5 min at room temperature, 50 μL of supernatant was transferred to a glass vial and subjected to GC-MS measurement.

### Gas chromatography coupled to mass spectrometry (GC-MS)

GC-MS analysis was performed using a GCMS-QP2010 Ultra (Shimadzu, Kyoto, Japan). The derivatized metabolites were separated on a DB-5 column (30 m x 0.25 mm id, film thickness 1.0 μm) (Agilent Technologies, Palo Alto, CA). Helium was used as the carrier gas at the flow rate of 39 cm/sec. The inlet temperature was set on 280°C. The column temperature was first held at 80°C for 2 min, then raised at the rate of 15°C/min to 330°C, and held for 6 min. One micro litter of the sample was injected into the GC-MS in the split mode (sprit ratio 1:3). The mass condition was as follows: electron ionization mode with the ionization voltage: 70 eV, ion source temperature: 200°C, interface temperature: 250°C, full scan mode in the range of m/z 85- 500, scan rate: 0.3 sec/scan. Data acquisition and peak processing were performed with a GCMS solution software Version 2.71 (Shimadzu, Kyoto, Japan).

### Semi targeted metabolomic data analysis

Chromatographic peaks were identified by comparing their mass spectral pattern to those in the NIST library or Shimadzu GC/MS Metabolite Database Ver. 1. The identification of metabolites was further confirmed through the coincidence of retention indices in samples with those in the corresponding authentic standards. Retention indices were determined and calibrated daily by the measurement of n-Alkane mixture (C8-40, Restek, USA), which was run at the beginning of batch analysis. A total of 101 metabolites were identified. Quantitative ions were selected for area calculation (summarized in supplementary Table 1) and the peak area of each metabolite was normalized.

### Animal experiments

C57BL/6J male mice were received from a commercial supplier (Janvier Labs, Courtaboeuf, France). A colony of Goto-Kakizaki (GK/Ox) rats was bred locally. Animals were maintained in specific pathogen free (SPF) condition. They had free access to a standard chow diet (R04-40, Safe, Augy, France) and were kept on 12h light/dark cycle. A group of 12 six-week old mice was fed control carbohydrate diet (CHD) (D 12450Ki, Research diets, NJ) and a group of 12 mice was fed high fat (60% fat and sucrose) diet (HFD) (D12492i, Research diets, NJ). One week later mice were anaesthetized with isoflurane and osmotic minipumps Alzet^®^ model 2006 (Charles River Lab France, l’Arbresle, France) filled with the bacterial metabolites 4-cresol or 4-methylcatechol (6 CHD-fed and 6 HFD-fed mice) (5.55mM in 0.9% NaCl) (Sigma Aldrich, St Quentin, France) or with saline (6 CHD-fed and 6 HFD-fed mice) were inserted subcutaneously on the dorsal left side. Concentration of 4-cresol was adjusted for a flow rate of 0.15μL/h. The same procedure was applied to insert the minipumps filled with 4-cresol (5.55mM in 0.9% NaCl) of saline subcutaneously in 32 week-old male GK rats.

Blood glucose and body weight were monitored weekly during the 6-week long administration of 4-cresol or saline. After three (mice) or four (rats) weeks of treatment (i.e. four weeks of HFD feeding in mice) an intraperitoneal glucose tolerance test (IPGTT) (2g/kg in mice, 1g/kg in rats) was performed in conscious animals following an overnight fast. For the IPGTT carried out in mice, blood was collected from the tail vein before glucose injection and 15, 30, 60 and 120 minutes afterwards. Additional blood samples were collected during the IPGTT in GK rats 15, 90, 180 and 240 minutes after glucose injection. Blood glucose levels were determined using an Accu-Check^®^ Performa (Roche Diagnostics, Meylan, France). Additional blood samples were collected at baseline and 30 minutes after glucose injection in Microvette^®^ CB 300 Lithium Heparin (Sarstedt, Marnay, France). Plasma was separated by centrifugation and stored at -80°C until insulin radioimmunoassay. Evaluation of overall glucose tolerance was obtained from the cumulative glycemia (GCum) and the ΔG, which were determined by the total increment of plasma glucose values during the IPGTT (GCum) and the cumulative glycemia during the test above baseline (ΔG).

After six weeks of metabolite treatment, animals were killed by decapitation and organs were dissected and weighed. Half of liver, fat and pancreas samples were snap frozen in liquid nitrogen and stored at -80°C for molecular studies, and the second half processed for histopathology.

All procedures were carried out under national French licence condition (Ref 00486.02) and were authorized following review by the institutional ethics committee of the University Pierre and Marie Curie.

### *In vitro* insulin secretion in mouse isolated islets

Eleven-weeks-old male C57Bl/6J (Janvier-Labs, Saint-Berthevin, France) were killed by cervical dislocation. A solution of collagenase (0.65mg/ml) (Roche, Sigma Aldrich, St Quentin, France) dissolved in a buffer containing Hanks (HBSS) (Gibco Invitrogen, France), Hepes (0.01M) (Gibco Invitrogen, France) and DNase I (0.1mg/ml) (Sigma Aldrich, St Quentin, France) was injected through the bile duct. Pancreas was dissected and incubated at 37°C in a collagenase solution. The reaction was stopped by addition of a cold buffer containing HBSS, BSA (0.5%) (Interchim, Montluçon, France) and Hepes (0.024M). Islets were purified on a four layer density gradient of Histopaque 1119 (Sigma-Aldrich, France) and HBSS, and incubated overnight at 37°C in RPMI (Gibco, Invitrogen, France) mixed with Hepes (10mM), sodium pyruvate (1mM) (Gibco Invitrogen, France), beta-mercaptoethanol (0.05mM), gentamicin (0.1mg/ml) (Sigma Aldrich, St Quentin, France) and FBS 10% (Eurobio, Les Ulis, France). Islets were then split into three groups incubated for 24 hours with either the culture medium (controls) or with cresol (10nM or 100nM) (W233706, Sigma Aldrich, St Quentin, France). *In vitro* insulin secretion tests were performed after 48 hours of cresol treatment. Islets were pre-incubated in KRBH-0.05% BSA with 2.8mM of glucose for 30 min, followed by 60 min incubation in KRBH-0.05% BSA with 2.8 or 16.7mM glucose to measure glucose-induced insulin secretion. Insulin was determined on supernatant by ELISA (Eurobio, Courtaboeuf, France). A batch of islets was used to determine insulin content.

### Histology and immunohistochemistry of animal tissues

Tissues were drop-fixed in 4% paraformaldehyde (Sigma-Aldrich, Saint Quentin Fallavier, France) immediately after collection and put through an automated carousel processor for dehydration, clearing, and paraffin embedding (Leica, Nanterre, France). Sections were prepared for liver (6 μm), pancreas (6 μm) and adipose tissue (10 μm), mounted on slides (DPX polymerizing mountant, Sigma-Aldrich, Saint Quentin Fallavier, France) and stained in Hematoxylin and Eosin (H&E). Epitope-specific antibodies were used for immunohistochemistry detection of insulin on pancreas sections (Dako, Saint Aubin, France).

For Oil red O (ORO), livers were snap frozen in OCT (VWR, Fontenay-sous-Bois, France) and cut into 7-μm sections using a cryostat. Sections were rehydrated in PBS (Sigma-Aldrich, St Quentin, France) and incubated with an ORO staining solution (Sigma Aldrich, St Quentin, France). Slides were washed in deionized water and mounted with Vectashield mounting medium (Laboratoires Eurobio Abcys, Les Ulis, France).

For immunohistochemistry analysis, pancreas sections were quenched with 3% H_2_O_2_, washed with TBS + 0.1 % (v/v) Tween-20 (or 0.05 % v/v Triton X-100 for nuclear epitopes), blocked with TBS + 3 % (w/v) BSA and incubated with diluted primary antibodies and then with HRP-conjugated secondary antibody (Bio-Rad, Marnes-la-Coquette, France). Chromogenic detection was carried out with the DAB chromogen kit (Dako, Saint Aubin, France). Nuclei were counterstained with hematoxylin. Quantitative expression of all immunostainings was performed using positive pixels algorithm (Indica Labs, Corrales, NM). For double immunostaining and immunofluorescence analyses, pancreas sections were stained for insulin as described above and co-stained with i) primary Anti-Ki 67 antibody (ab15580, Abcam, Paris, France) and secondary Donkey Anti-Goat IgG H&L conjugated to Alexa Fluor^®^ 568 (ab175704, Abcam, Paris, France) ii) primary Mouse/Rat CD31/PECAM-1 Antibody (AF3628, Minneapolis, USA) and secondary Goat anti-Rabbit IgG H&L conjugated to Alexa Fluor 488 (A-11034, ThermoFisher, Villebon, France). Results are expressed as percentage of positive pixels, within islets where indicated. The quantification method is an automated observer-independent process based on section scanning and application of publicly available algorithms. Each biological replicate represents one slide per animal mounted with at least 3 tissue sections, representing 3 technical replicates, the mean and variance of which is presented as the result per biological replicate. All images were acquired on an Axiovert 200M microscope (Zeiss, Marly-le-Roi, France).

### Histology of mouse islets of Langerhans

Islet immunohistochemistry was carried out after 48 hours of cresol treatment on 100 islets washed with DPBS (Gibco, Invitrogen, France), fixed in paraformaldehyde 4% (Sigma Aldrich, St Quentin, France), washed again and centrifuged. A solution of preheated agarose (10 mg/ml) (Eurobio Abcys, Les Ulis, France) containing heamatoxylin was added on the pellets. Cooled agar gels containing islets were picked and transferred in ethanol 70% at 4°C. Islets were dehydrated and embedded in paraffin. Islets sections (5 μm) were prepared and quenched with 3% H_2_O_2_, washed with TBS +0.1% (v/v) Triton X-100 (Sigma-Aldrich, St Quentin, France), blocked with TBS +3 % (w/v) BSA and incubated with diluted primary antibodies against Ki67 (ab16667, Abcam, Paris, France) and then with HRP-conjugated secondary antibody (Bio-Rad, Marnes-la-Coquette, France). Chromogenic detection was carried out with the DAB chromogen kit (Dako, Saint Aubin, France). Nuclei were counterstained with hematoxylin. Positive and negative nuclei were counted on all islets. Results are expressed as percentage of positive Ki67 cells. We counted 25 to 33 islets for each cresol concentration. All images were acquired on an Leica DM4000 B microscope (Leica, Nanterre, France).

### RNA isolation and quantitative RT-PCR

RNA was extracted from pancreas and adipose tissue using the RNeasy RNA Mini Kit (Qiagen, Courtaboeuf, France). Reverse transcription was performed from a 20 μL reaction mixture with 500 ng RNA using M-MLV reverse transcriptase kit (ThermoFisher, Villebon, France). Quantitative RT-PCR was performed using sequence specific primers and the MESA green kit for SYBR green assays (Eurogentec, Angers, France). We used 18S and/or cyclophilin housekeeping genes to normalize relative quantification of mRNA levels using the Livak and Scmittgen methods (*2*). Primers are listed in Supplementary Table 2.

### Analytical assays

Blood glucose was measured using an Accu-Check^®^ Performa (Roche Diagnostics, Meylan, France) and plasma insulin was determined using Insulin ELISA kits (Mercodia, Uppsala, Sweden). For determination of liver triglycerides, liver samples (100mg) were homogenised and incubated in Nonidet P-40 (5%) and supernatants containing triglycerides were collected. Triglycerides concentration was quantified in the supernatant fraction using a colorimetric assay (ab65336, Abcam, Paris, France) by measuring OD at 570nm. The ratio NAD/NADH was determined using a quantification kit (MAK037; Sigma Aldrich, St Quentin, France) on extracts prepared from 20 mg pancreas tissue. Samples were homogenized in the extraction buffer and clarified by centrifugation. Supernatant was deproteinized by filtration through a 10-kDa cutoff spin filter (Millipore SAS, Molsheim, France). The assay was then performed according to the manufacturer’s instructions.

### Statistical analyses

Statistical analysis of targeted metabolomic data was performed to assess association of each metabolite with CMD risk variables (diabetes, hypertension, hyperlipidemia, body weight and body mass index) based on a linear regression model. Normality assumption of the residuals of each metabolic feature was investigated by Shapiro-Wilk test. The R statistical language was then used to perform the linear regression and compute a P-value for each metabolic feature with a threshold of significance set to 0.05. False discovery rates (FDR) were corrected using the Benjamini-Hochberg method to adjust P-values for false discovery involving multiple comparisons. Analyses were adjusted for age and gender effects. Statistical analyses of results from animal studies were performed using the GraphPad Prism 6 software. Statistics were calculated using two-way ANOVA, two-tailed Student t test or Mann Whitney test. Differences were considered statistically significant with a P<0.05.

Multivariate modelling of the signature related to diabetes and obesity was performed through Orthogonal Partial Least Squares Discriminant Analysis (O-PLS-DA) (*3*). The O-PLS-DA models were validated by 7–fold cross-validation. The significance of model goodness-of-fit (R^2^_Yhat_) and 7-fold CV goodness-of-prediction (Q^2^_Yhat_) parameters was tested by resampling 100,000 times the model under the null hypothesis (i.e. randomly permuted class membership, not related to clinical diagnosis) as previously described (*4*). To confirm O-PLS-DA model coefficients, we fitted a logistic regression model adjusted for age and gender for each metabolite. The class membership/outcome variable was coded as follows: (0:non-obese, non-diabetic; 1: obese and/or diabetic). All p-values were corrected for multiple testing using the Benjamini-Hochberg procedure.

